# NUSAP1 regulates mitotic processes via KIF2C interaction and AURKA phosphorylation in primary microcephaly

**DOI:** 10.1101/2025.05.16.654427

**Authors:** Decheng Ren, Keyi Li, Zhen Liu, Yan Bi, Liangjie Liu, Lei Ji, Ke Yang, Yingying Luo, Le Luo, Yang Yan, Yang Li, Fengping Yang, Hua Wang, Guang He, Xiao Mao

**Author notes:** These authors contributed equally to this work (Decheng Ren, Keyi Li, Zhen Liu). Corresponding author at: National Health Commission Key Laboratory for Birth Defect Research and Prevention, Hunan Provincial Maternal and Child Health Care Hospital, Changsha 410100, China.; Bio-X Institutes, Shanghai Jiao Tong University, 1954 Huashan Road, Shanghai 200030, China.

## Abstract

Primary microcephaly (PM) is a neurodevelopmental disorder characterized by a significantly smaller head than normal. Despite the identification of several genes associated with PM subtypes, the etiology remains unclear in a significant proportion of patients. Here, we reported two de novo nonsense mutations in the *NUSAP1* gene identified in two independent PM families: c.1209 C>G (p.Tyr403*) and c.1213C>T (p.Gln405*). *Nusap1*-edited mouse models, designed to mimic Human *NUSAP1* mutations, replicated the microcephaly phenotype, characterized by a small brain and thinner cerebral cortex. We demonstrated that *Nusap1* was specifically expressed in neural stem/progenitor cells (NSPCs), and the number of NSPCs was significantly reduced in *Nusap1*-edited mice. In vitro experiments revealed that *NUSAP1* mutations disrupted mitotic metaphase chromosome alignment, increased polyploidy, induced cell cycle arrest, and promoted apoptosis. Co-IP confirmed direct interactions between NUSAP1 and KIF2C, and showed that truncating mutations impaired their binding. Furthermore, NUSAP1 rescued the cell cycle arrest and increased apoptosis caused by KIF2C overexpression in *NUSAP1* KO HEK293T cells, whereas the mutant truncated protein failed to exert such a rescue effect. We also uncovered an interaction between NUSAP1 and Aurora kinase A (AURKA). Intriguingly, subsequent experiments revealed that AURKA-mediated phosphorylation of NUSAP1 modulates its binding affinity to KIF2C. In conclusion, our findings established *NUSAP1* as a novel pathogenic gene for PM. We proposed a model in which AURKA-mediated phosphorylation of NUSAP1 regulates its interaction with KIF2C, balancing spindle microtubule stability and depolymerization to ensure proper sister chromatid segregation. Disruptions in this pathway impair NSPCs mitosis, thereby affecting neocortical development.

## Introduction

Primary microcephaly (PM) is a neurodevelopmental disorder characterized by a smaller-than-expected head circumference in fetuses or infants compared to peers of the same gestational age, sex, and ethnicity(1–3). The incidence of primary non-syndromal microcephaly is 1:30,000 to 1:250,000 live births(4). Microcephaly pathogenesis depends on genetic and environmental factors(5). Most primary microcephaly cases follow autosomal recessive inheritance, though autosomal dominant inheritance also occurs. Compared with the relatively clear gene-phenotype correspondence in autosomal recessive inheritance, cases of autosomal dominant inheritance exhibit significant and pronounced heterogeneity(6). To date, 30 autosomal recessive primary microcephaly (MCPH) genes have been identified, yet the causes remain unknown in many patients(7, 8). Pathogenic MCPH genes predominantly encode proteins localized to the mitotic apparatus, such as centrosomes and spindles(9). For instance, ASPM or CENPJ mutations disrupt spindle pole assembly and chromosome segregation during mitosis, reducing neural stem/progenitor cell (NSPC) pools and resulting in smaller brain size(10). Environmental factors like maternal teratogen exposure or Zika virus infection can further impair NSPC proliferation and differentiation(11–13). Despite research progress, the molecular mechanisms of microcephaly are still not fully understood, hindering the development of precise diagnostic and treatment strategies.

The mitotic spindle, a dynamic microtubule-based structure, plays a crucial role in ensuring accurate chromosome segregation during cell division(14). This function is especially vital in the developing brain, where proper spindle activity is indispensable for the proliferation, differentiation, and migration of NSPCs. However, spindle abnormalities, such as defects in microtubule polymerization, spindle pole organization, or motor protein function, can disrupt chromosome segregation. These errors can result in aneuploidy, apoptosis, or dysregulated cell division patterns in NSPCs(15, 16). During neurogenesis, these spindle-related defects in NSPCs disrupt the delicate balance between NSPCs self-renewal and differentiation. As a consequence, the production of neurons is ultimately reduced, which is a characteristic feature of microcephaly. Mutations in spindle-associated proteins like KIF2A could result in spindle assembly defects and impair neurogenesis in mouse models, effectively recapitulating microcephaly phenotypes(17). Moreover, research using human induced pluripotent stem cell (iPSC) - derived NSPCs has demonstrated that spindle dysfunction indeed derails neurodevelopmental processes(18). Collectively, these findings strongly emphasize the pivotal role of spindle dynamics in brain development and its significant association with the pathogenesis of microcephaly.

The *NUSAP1* (nucleolar and spindle - associated protein 1) gene encodes a conserved microtubule - associated protein (MAP) of approximately 55 kDa, which is selectively expressed in proliferating cells(19, 20). Mechanistically, NUSAP1 stabilizes microtubules through cross-linking, bundling and attachment to chromosomes, thereby ensuring proper spindle formation, chromosome alignment, and accurate segregation during cell division(21–23). Recent investigations in mouse oocytes have highlighted the critical importance of precise NUSAP1 regulation. Its dynamic expression and localization are essential for maintaining meiotic fidelity, supporting spindle integrity, and coordinating cytoplasmic organization during oocyte maturation(24). The protein levels of NUSAP1 are tightly regulated by the anaphase-promoting complex/cyclosome (APC/C) during the cell cycle, and high expression of *NUSAP1* was observed in several types of cancers(25). Although the roles of NUSAP1 in the division of tumor cells and oocytes have been studied, its functions in brain development and the occurrence of microcephaly remain poorly understood.

In this study, whole-exome sequencing (WES) identified two de novo nonsense mutations (c.1209C>G, c.1213C>T) in *NUSAP1* from two primary microcephaly (PM) families. We showed that NUSAP1 is phosphorylated by AURKA, and these mutations impaired its interaction with KIF2C, disrupting mitotic chromosome alignment, increasing polyploidy, arresting the cell cycle, and promoting apoptosis. Moreover, Nusap1-edited mice developed microcephaly, validating NUSAP1’s pathogenic role in vivo. Collectively, our findings reveal a novel mechanism by which NUSAP1 mutations dysregulate the mitotic spindle, cell cycle, and apoptosis via weakened interaction with KIF2C.

## Materials and Methods

### Patients and Genetic Analysis

The two patients were recruited from Hunan Provincial Maternal and Child Health Care Hospital. The parents of both patients were included for genetics analysis and provided informed consent. This study was approved by the Ethics Committee of Maternal and Child Health Hospital of Hunan Province (2025028).

For whole-exome sequencing, whole blood samples were collected with EDTA Blood Collection Tubes (Becton, Dickinson and Company, NJ, USA). DNA was extracted using FlexiGene DNA kits (Fuji, Tokyo, Japan) following the standard procedures. DNA libraries were prepared according to the instructions from Illumina (Illumina, San Diego, CA). Whole-exome enrichment was performed using the TruSeq Exome Enrichment Kit (Illumina, San Diego, CA) with the recommended protocol. Captured DNA libraries were sequenced with HiSeq2500 (Illumina, San Diego, CA, USA).

We firstly used cutadapt (v1.15) to trim adaptor sequences at the tail of sequencing reads, and then aligned sequencing reads to human reference genome (UCSC hg19) with BWA (v0.7.15). Duplicated reads were marked by Picard (v2.4.1). Qualimap (v2.2.1) was used to calculate base quality metrics, genome mapping rate, and the coverage of targeted regions. Base quality score recalibration, indel realignment and variants (SNVs & InDels) calling were performed following the best practice protocol of the Genome Analysis Toolkit (GATK, v3.8). Variant filtering was done by a finely tuned in house script. Pass-filter variants were annotated using Pubvar variant annotation engine (www.pubvar.com) and VEP (release 88). In genetic analysis, we separately identified variants that fit the dominant and recessive inheritance models. Variants met anyone of the following criteria were excluded from genetic analysis: maximum population frequency was large than 0.01, genotype confidence was low, or predicted as benign by SIFT and PolyPhen 2. The pathogenic evidences of candidate disease-causing variants were scored by InterVar (1.0.8) according to ACMG guidelines(26). All the above analysis was performed on Seqmax (www.seqmax.com).

### Plasmid Construction and Mutagenesis

Human *NUSAP1* (NM_016359.5) was amplified from cDNA using PrimeSTAR® GXL Premix (Takara, Dalian, China) and inserted into pEGFP-C3 or pCMV-HA-N vectors. Human *KIF2C* (NM_006845.4) was amplified from cDNA and inserted into pmCherry-N1 or p3xFlag CMV-14 vectors. Human *AURKA* (NM_198433.3) was also amplified from cDNA and inserted into p3xFlag CMV-14 vectors. DNA ligation reactions were executed using Solution I from the DNA Ligation Kit (Takara). The reactions were incubated at 16°C for a duration of 2 hours to ensure efficient vector - insert conjugation. Site-directed mutagenesis was conducted to generate mutant plasmids by using the Hieff Mut™ Site-Directed Mutagenesis Kit (YEASEN, Shanghai, China). Primers for PCR amplification and *NUSAP1* mutagenesis are listed below:

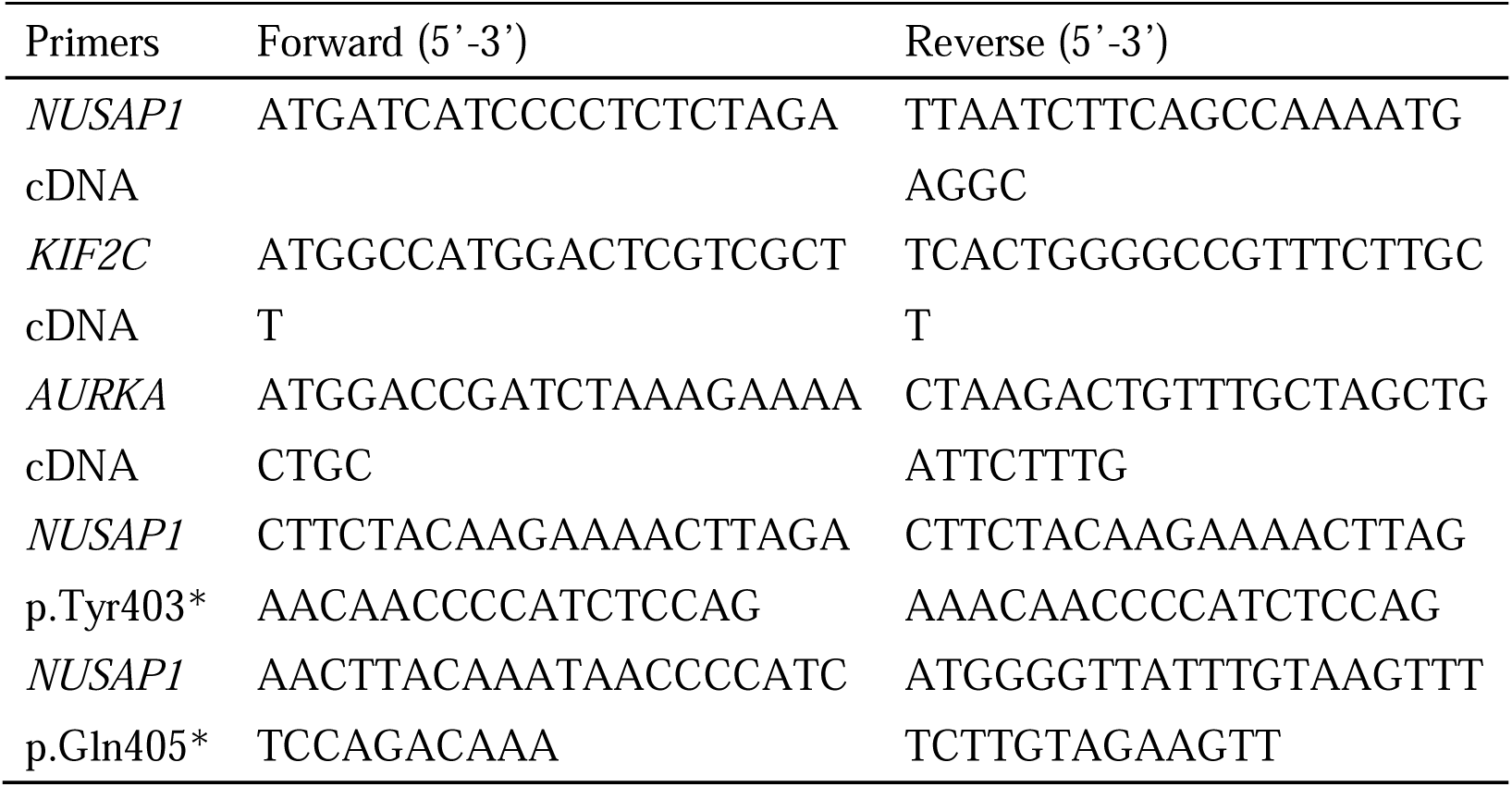

### Construction of Gene-edited Mice

The sgRNA was designed to target the sequence (sgRNA: GCGCACCTGGTTTGGAGATGAGG) of *Nusap1* (NM_133851.3) gene. The pCD-CAS plasmid (Biotechnologies) carrying the *Nusap1* sequence was used as the template for PCR with the primer-sgRNA and T7R1. The PCR products were purified and used for in vitro RNA synthesis with a MEGASHORT T7 high yield Transcription Kit (Ambion). The sgRNA was purified with a MEGAclear-96 Purification of Transcription Reactions Kit (Ambion). The pT7-cas9 plasmid (Biotechnologies) was linearized by Xba I (Takara) and transcribed into mRNA in vitro using a mMESSAGE mMACHINE™ T7 ULTRA Transcription Kit (Ambion). Approximately 5 pL of the mixture of the Cas9 mRNA (20 ng/μL) and sgRNA (10 ng/μL) was injected into the cytoplasm of C57BL/6J zygote with a 2-3 μm diameter pipette using a Piezo instrument (PIEZOXPERT, Eppendorf). After microinjection, zygotes were cultured in KSOM medium in a humidified atmosphere containing 5% CO_2_ and 95% air at 37°C for 24 h. The embryos at the 2-cell stage were surgically transferred to both the oviducts of recipient ICR mice. Genomic DNA was extracted from the toe tissues of one-week old mice by using a TIANGEN Genomic DNA kit (TIANGEN). The fragments were amplified from genomic DNA by PCR using the primers (primer forward: GGCAGGCTGAAAGAT; primer reverse: AGGCTGTTGGGAAGA). Six F0 mice were born and a messy peak appeared near the sgRNA in one of the mouse sequencing results. F1 mice were further obtained and one mouse line carrying 13bp deletion was subjected to the current studies.

### In Utero Electroporation (IUE)

IUE was performed according to previously published protocol(27). The human *NUSAP1* gene or the *NUSAP1* gene carrying the p.Gln405* mutation was cloned into the pCAGIG plasmid. This plasmid, a commonly - used vector in in utero electroporation (IUE), contains an IRES2 sequence that allows for relatively independent expression of the target gene and GFP. Plasmid was prepared using EndoLJFree PlasTmid Maxi Kit (QIAGEN) and diluted to 2∼3 μg/μl. Plasmid was electroporated into the dorsal cortex of the E14.5 embryonic mouse brain using ECM830 Electroporation system (BTX).

### *NUSAP1* Knock Out (KO) Cells

The *NUSAP1* polyclonal knockout (KO) cells were established based on HEK293T using CRISPRLJCas9 gene editing technique. Plasmids were transfected into cells by nucleofection using Neon Transfection system (Invitrogen) according to the manufacturer’s guidance. The sequence of sgRNA is sgRNA-*NUSAP1*: GGAGTCCAGCTCCTCTAGAG. *NUSAP1* KO cells simultaneously expresses Cas9, the sgRNA and the puromycin resistance gene. After being screened with puromycin, the protein expression was detected by Western blot.

### Cell Culture and Transfection

HEK-293T cells were cultured in DMEM (HyClone) supplemented with 10% FBS (Gibco) at 37°C in a 5% COLJ atmosphere. Transient transfection was performed using FuGENE® HD reagent (Promega) according to the manufacturer’s instructions. Cells were seeded at 5×10LJ cells per well in 6-well plates. After 24 hours, 3.3 µg of plasmid DNA diluted in 155 µl Opti-MEM (Gibco) was mixed with 9.9 µl FuGENE® and added to the cells.

### Immunofluorescence (IF) and Immunocytochemistry (ICC) Microscopy

Mouse brains were fixed in 4% paraformaldehyde (PFA) in PBS at 4°C overnight. Then, tissues were transfer to 30% sucrose overnight for dehydration and embedded in optimal cutting temperature compound (OCT) (Sakura). Mouse brains were cryosectioned at 18 μm (Leica CM3050S). The organoid sections of human embryonic stem cells (hESC) were kindly donated by Zhenming Guo from the School of life sciences and technology, Tongji University. For antigen retrieval, brain sections were buffered in boiling antigen recovery solution (1 mM EDTA, 5 mM Tris at pH 8.0) for 15 min and cooled to RT. Sections were blocked at RT for 1 h in 0.5% Trion XLJ100 and 4% bovine serum albumin (BSA). Sections were incubated with primary antibody at 4°C overnight and then incubated with secondary antibody at RT for 1 h.

Cells grown on coverslips were fixed in precooled methanol for 5 minutes, blocked with 3% BSA, and incubated with primary antibodies at 4°C overnight. After washing, sections or cells were stained with fluorescence-conjugated secondary antibodies (Invitrogen) for 1 hour at room temperature. Coverslips were mounted with Mowiol (Sigma-Aldrich) containing 1 µg/ml DAPI. Images were acquired using confocal microscopes (Leica TCS SP8) with 63×/1.4 NA oil objectives.

Primary antibodies used in this study for immunofluorescence were anti-phospho-Histone H3 (#53348, Cell Signaling Technology, 1:500); anti-Ki67 (#9129, Cell Signaling Thechnology, 1:500); anti-NUSAP1(12024-1-AP, Proteintech,1:500); anti-SOX2(MA1-014, ThermoFisher, 1:200), anti-PAX6 (42-6600, ThermoFisher, 1:200); anti-KIF2C (MA5-25647, ThermoFisher, 1:200); anti-AURKA (45-8900, ThermoFisher, 1:200); anti-α-Tubulin (ab7291, Abcam, 1:500); anti-γ-Tubulin (ab179503, Abcam, 1:500); anti-BUBR1 (ab209998, Abcam,1:500); DAPI (D9542, Sigma-Aldrich). The following secondary antibodies were used: Alexa Fluor 488-conjugated anti-mouse IgG (H+L; A-21202; Invitrogen), Alexa Fluor 488-conjugated anti-rabbit IgG (H+L; A-21206; In-vitrogen), Alexa Fluor 594-conjugated anti-mouse IgG (H+L; A-21203; Invitrogen), Alexa Fluor 594-conjugated anti-rabbit IgG (H+L; A-21207; Invitrogen), Alexa Fluor 647-conjugated anti-rabbit IgG (H+L; A-31573; Invitrogen).

### Western blot (WB) Analysis

HEK-293T cells were lysed in RIPA buffer (Sigma). Total lysates were mixed with SDS-PAGE loading buffer (Beyotime), denatured by boiling for 10 minutes, and separated on 10% polyacrylamide gels. Proteins were transferred to PVDF membranes (GE Healthcare), blocked with Western blot quick blocking buffer (Beyotime), and incubated overnight with primary antibodies at 4°C. HRP-conjugated secondary antibodies (Jackson ImmunoResearch, 1:5000) were applied for 1 hour at room temperature. Blots were developed using ECL reagent (Share-bio).

Primary antibodies used in this study for WB were anti-GFP (ab290, Abcam, 1:5000), and anti-NUSAP1(12024-1-AP, Proteintech,1:5000); anti-Flag (30505ES60, Yeasen, 1:5000); anti-HA (30704ES60, Yeasen, 1:5000); anti-GAPDH (HC301-01, TransGen,1:5000). HRP-conjugated anti-rabbit IgG (323-005-021; Jackson), and HRP-conjugated anti-mouse IgG (223-005-024; Jackson)

### Co-Immunoprecipitation (Co-IP) Analysis

For Co-IP of GFP-KIF2C/HA-NUSAP1, HEK293T cells transfected with HA/HA-NUSAP1 and GFP/GFP-KIF2C were lysed on ice for 20 minutes in an ice-cold NP40 lysis buffer (Beyotime). The lysates were then incubated with anti-GFP or anti-HA nanobody magarose beads (AlpaLifeBio) for 1 h at 4°C.

For Co-IP of Flag-AURKA/HA-NUSAP1 or Flag-KIF2C/HA-NUSAP1, HEK293T cells transfected with HA/HA-NUSAP1 and Flag/Flag-AURKA or Flag/Flag-KIF2C were lysed. The lysates were incubated with anti-HA or anti-Flag nanobody magarose beads (AlpaLifeBio) for 1 h at 4°C as well.

After washing the beads five times with PBST buffer, loading buffer was added to resuspend the beads. Then, the beads were incubated in a 95°C water bath for 10 minutes to dissociate the immunoprecipitation complexes from the beads. Subsequently, the magnetic beads were separated from the solution by placing the tube on a magnetic rack and allowing it to stand for a short period. Finally, the samples were subjected to Western blotting analysis to detect the target proteins.

### Terminal deoxynucleotidyl transferase dUTP nick end labeling (TUNEL) assay & microscopy

TUNEL assay was performed according to manufacturer’s instructions (One Step TUNEL Apoptosis Assay Kit, C1090, Beyotime) to detect cell death in the mouse cortex. The assay stained in the red channel at 568 nm. DAPI was applied as a nuclear counter-stain in the blue channel at 461 nm. Images were acquired using confocal microscopes (Leica TCS SP8) with 20×objectives. Exposure settings were adjusted to minimize oversaturation.

### 5-Ethynyl-2′-deoxyuridine (EdU) Assay for Cell Proliferation

Cell proliferation was assessed using the BeyoClick™ EdU-594 Cell Proliferation Kit (Beyotime). Transfected cells were incubated with EdU for 2 hours, fixed, and stained according to the manufacturer’s protocol. EdU-positive cells were visualized by fluorescence microscopy (Axio Imager M2, Zeiss) and quantified using ImageJ software. The proliferation index was calculated as (EdU-positive cells/DAPI-positive cells) × 100%.

### Cell Cycle Analysis

For the cell cycle analysis, after transfection, the cells were harvested from each well by trypsinization and fixed in 70% (v/v) cold ethanol at 4 °C overnight. After washing with ice-cold PBS, the fixed-cell pellets were collected by centrifugation. Cells were prepared by using Cell Cycle and Apoptosis Analysis Kit (Beyotime) following procedures and resuspended in Propidium/RNase Staining Buffer for staining of DNA and finally analyzed using flow cytometry (Beckman Cytoflex).

### Detection of Apoptosis

To quantify cell apoptosis, after the indicated transfections, the cells were collected, and the apoptotic portion was identified using the Annexin V-FITC/PI Apoptosis Detection Kit (Beyotime) following the manufacturer’s instructions. The number of apoptotic cells was then quantified by flow cytometry (Beckman Cytoflex), and the analysis was carried out using FlowJo software (version 10.5.2).

### Quantitative PCR (qPCR)

Total RNA was extracted using TRIzol reagent (Tiangen), and cDNA was synthesized with the Hifair® II 1st Strand cDNA Synthesis Kit (YEASEN). qPCR was performed on a LightCycler® 480 Instrument II (Roche) usingTB Green® Premix Ex Taq™ II (Tli RNaseH Plus) (Takara). Primers are listed below, with GAPDH as the internal control.

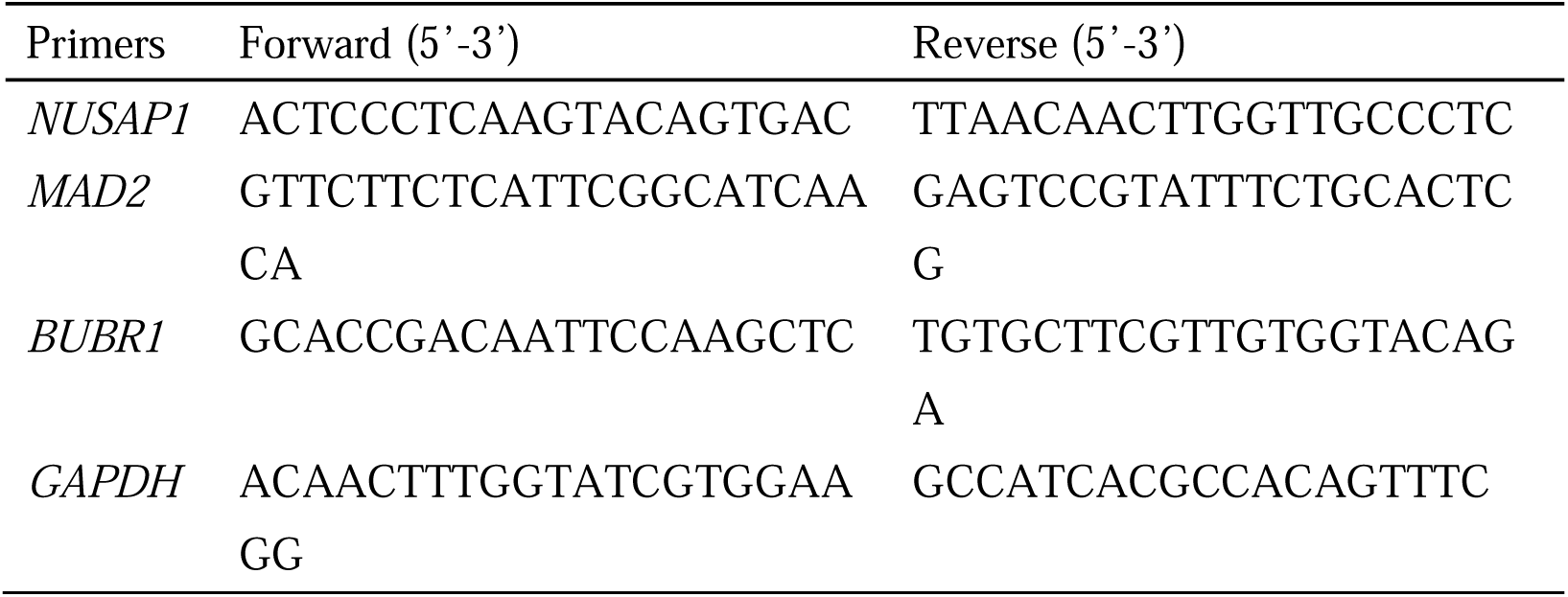

### Statistical Analysis

Data were analyzed using GraphPad Prism 8.4.3 (GraphPad Software). Normality was assessed by the Kolmogorov-Smirnov test, and group differences were evaluated using the independent two-tailed t-test. Results are presented as mean±SEM, with statistical significance set at P < 0.05. Significant difference was marked with asterisks (*, *P*<0.05; **, *P*<0.01; ***, *P*<0.001).

## Results

### Clinical Presentation of Affected Individuals

We performed the systematic trios-based WES in two independent PM families and identified two de novo nonsense mutations in *NUSAP1* (c.1213C>T (p.Gln405* and c.1209 C>G (p.Tyr403*)). Both mutations were confirmed by Sanger sequencing (Fig. 1B, C). Both mutations can lead to the truncation of the NUSAP1 protein, resulting in the loss of two domains, namely the ChHD domain and MCBD domain (Fig. 1D, E).

**Figure 1.**
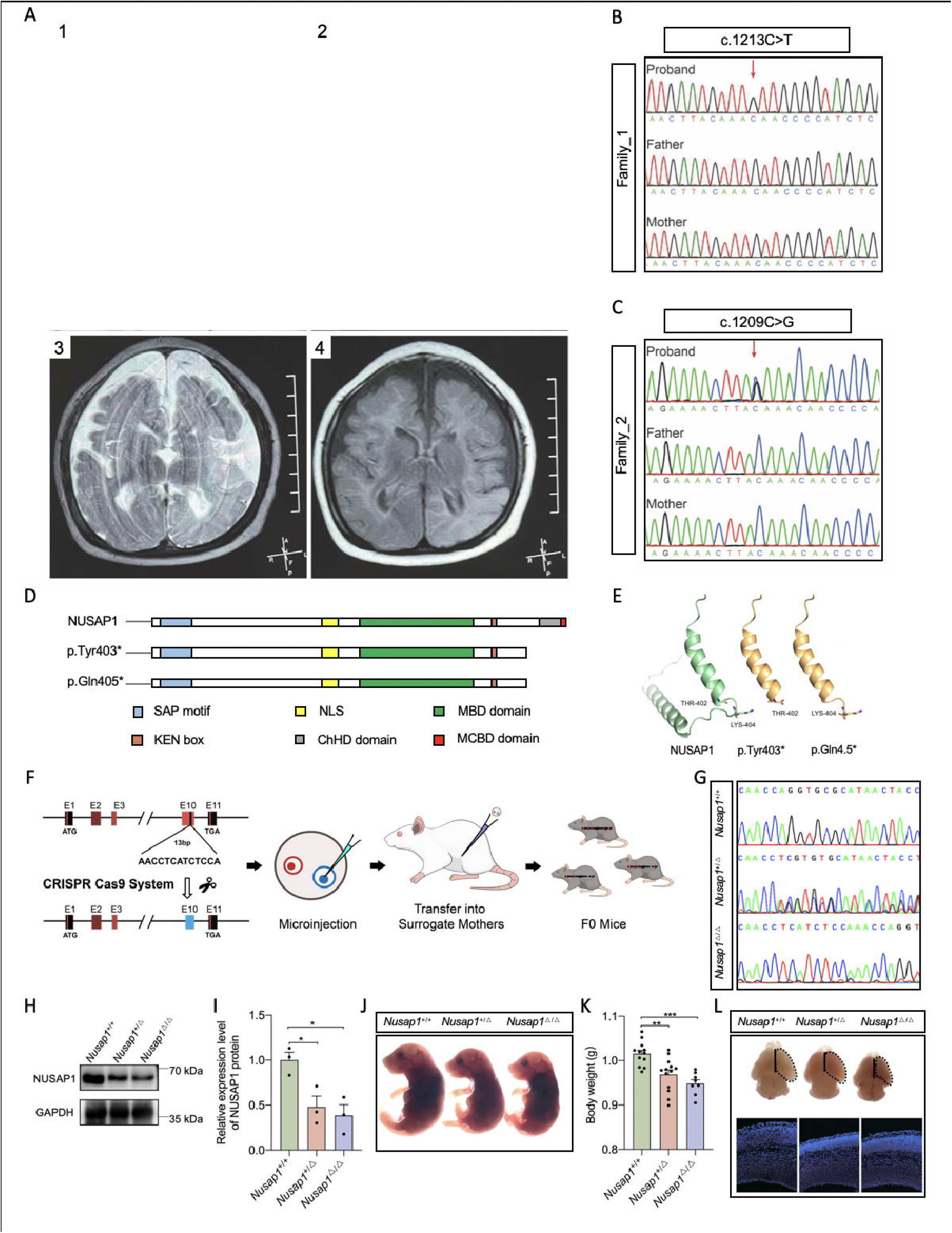
Clinical, genetic, and phenotypic insights into NUSAP1 mutations in humans and mice. **(A)** Photographs of Family 1 proband at 2 months (A-1) and 7 years 5 months (A-2), showing microcephaly. (A-3,4) Brain MRI at 2 years of age (Family 1). **(B)** Sanger sequencing chromatogram of Family 1, revealing a heterozygous nonsense variant in *NUSAP1* (c.1213C>T, p.Gln405*). **(C)** Sanger sequencing of Family 2, showing a heterozygous nonsense variant in *NUSAP1* (c.1209C>G, p.Tyr403*). **(D)** The schematic diagram shows the secondary structure of the protein. Different colors are used to label the protein domains. Both mutations lead to the truncation of the amino acid sequence and the loss of ChHD domain and MCBD domain. **(E)** Protein structure simulation shows that the NUSAP1 mutation results in the loss of the terminal structure of the protein. **(F)** The experimental schematic diagram of *Nusap1* gene editing. **(G)** The genotypes of mice were accurately determined via Sanger sequencing. **(H)** Western blot analysis was performed to detect the NUSAP1 expression in the mice cerebral cortex. **(I)** Quantification of the protein expression levels in three groups. **(J)** At embryonic day 14.5 (E14.5), WT mice were larger in body size than their heterozygous and homozygous littermates. **(K)** Quantification of the body weight levels in three groups. **(L)** At E14.5, both the cerebral cortex size and the cortical thickness in heterozygous and homozygous mice were smaller and thinner, respectively, compared to those in their WT littermates.

The proband in family_1 was a 7-year 5-month-old male with severe global developmental delay, microcephaly (Fig. 1A1, 2), and drug-resistant epilepsy. Born to nonconsanguineous Chinese parents with no family medical history, microcephaly was detected prenatally (21.2 cm at 26 weeks, 24.7 cm at 30 weeks, 28.6 cm at 36 weeks, all < -3SD). Term delivery was complicated by amniotic fluid aspiration-induced asphyxia (Apgar scores 8, 9, 9 at 1, 5, 10 minutes), requiring 9 days of intubation and mechanical ventilation. His birth head circumference was 30 cm (-3.5SD). Right ocular clonic seizures started on day 17, with EEG showing interictal right-sided sharp waves and ictal spike-slow wave discharges. Despite treatment with levetiracetam, sodium valproate, and clonazepam, he had multiple seizure types lasting 10 seconds to 30 minutes, accompanied by postictal sleepiness, consciousness impairment, and feeding refusal. Brain MRI at 2 years revealed partial corpus callosum agenesis (Fig. 1A3, 4). At the latest follow-up, daily seizures persisted. His head circumference was 44 cm (< -6SD), while height (130 cm) and weight (28 kg) were normal for age. Severely developmentally delayed, he only achieved basic motor skills like unsteady head lifting and rolling, lacked speech, and required a liquid diet due to dysphagia. Neurologic examination showed generalized hypertonia and precocious puberty signs. All of the above phenotypic features can be found summarized in Table S1.Trio-WES identified a novel heterozygous nonsense variant (NM_016359.4: c.1213C>T, p.Gln405*) in *NUSAP1* exon 10, absent from public databases.

**Figure 2.**
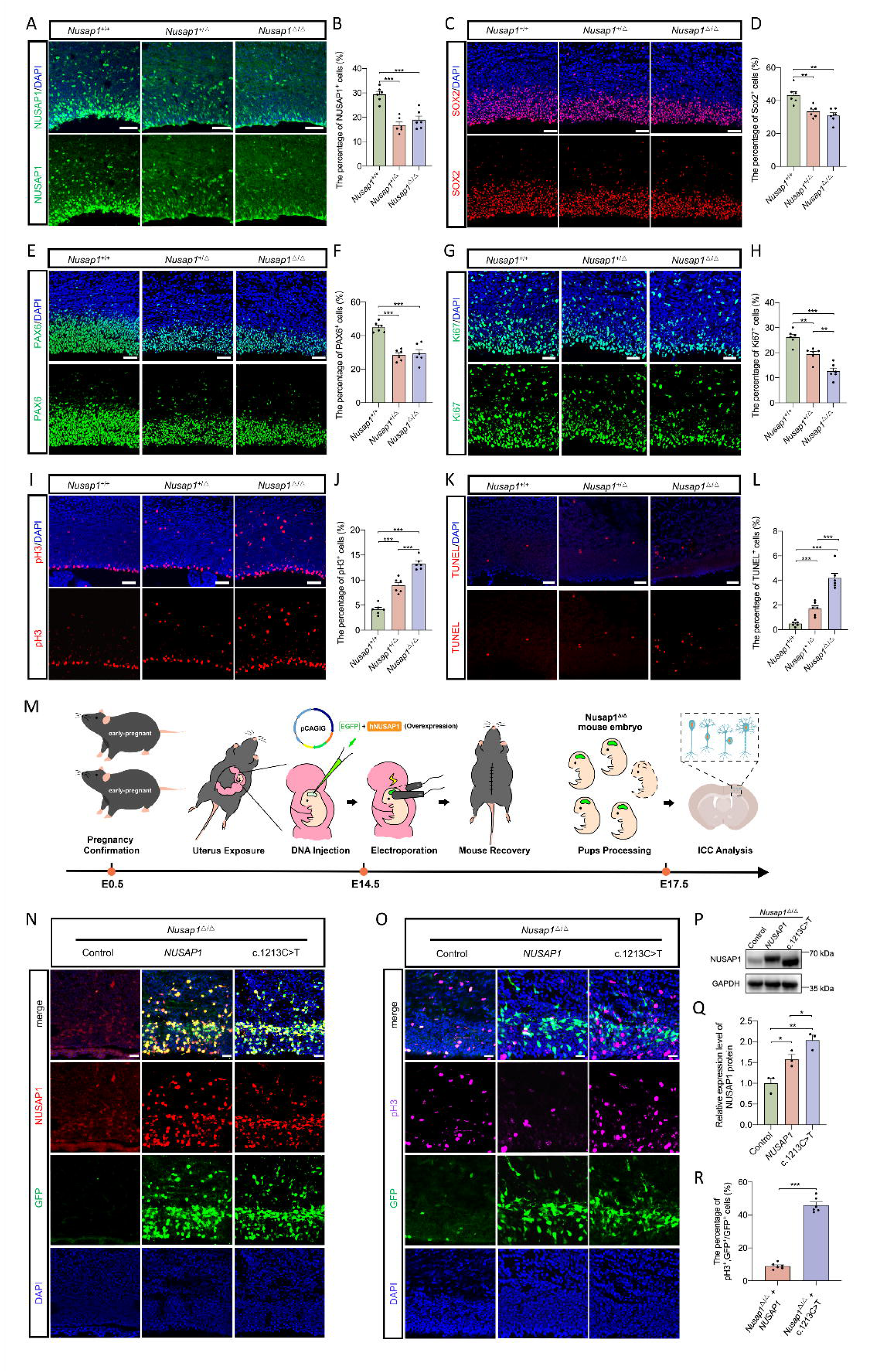
Immunofluorescent analyses and functional assays of NUSAP1 in cerebral cortex development. **(A)** Immunofluorescent images of the cerebral cortex stained for NUSAP1 (green) to detect NUSAP1 positive cells. The merged images of the NUSAP1 and DAPI channels were shown above. Scale bar = 45 μm. **(B)** Quantification of the percentage of NUSAP1-positive cells in the cerebral cortex. **(C)** Immunofluorescent images of the cerebral cortex stained for SOX2 (red) to detect NSPCs. The merged images of the SOX2 and DAPI channels were shown above. Scale bar = 45 μm. **(D)** Quantification of the percentage of SOX2-positive cells in the cerebral cortex. **(E)** Immunofluorescent images of the cerebral cortex stained for PAX6 (green) to detect NSPCs. The merged images of the PAX6 and DAPI channels were shown above. Scale bar = 45 μm. **(F)** Quantification of the percentage of PAX6-positive cells in the cerebral cortex. **(G)** Immunofluorescent images of the cerebral cortex stained for Ki67 (green) to detect proliferating cells. The merged images of the Ki67 and DAPI channels were shown above. Scale bar = 45 μm. **(H)** Quantification of the percentage of Ki67-positive cells in the cerebral cortex. **(I)** Immunofluorescent images of the cerebral cortex stained for pH3 (red) to detect cell cycle arrested cells. The merged images of the pH3 and DAPI channels were shown above. Scale bar = 45 μm. **(J)** Quantification of the percentage of pH3-positive cells in the cerebral cortex. **(K)** Immunofluorescent images of the cerebral cortex stained for TUNEL (red) to detect apoptotic cells. The merged images of the TUNEL and DAPI channels were shown above. Scale bar = 50 μm. **(L)** Quantification of the percentage of TUNEL-positive cells in the cerebral cortex. **(M)** The experimental schematic diagram of in utero electroporation. Human derived *NUSAP1* and c.1213C>T plasmids were electroporated into the cortical cells of E14.5 *Nusap1*^△/△^ mice. Mice were sacrificed and brain tissues were harvested three days later. **(N)** Immunofluorescent images of the cerebral cortex stained for NUSAP1 (red) to detect the expression of exogenous plasmids. GFP represents the plasmid-electroporated-positive cells. Merged images of the NUSAP1, GFP, and DAPI channels were shown above. Scale bar = 50 μm. **(P)** Western blot analysis was performed to detect the NUSAP1 expression in the cerebral cortex. **(Q)** Quantification of the protein expression levels in three groups. (**O)** Immunofluorescent images of the cerebral cortex stained for pH3 (purple) to detect mitotic cells. GFP represents the plasmid-electroporated-positive cells. The merged images of the pH3, GFP, and DAPI channels were shown above. Scale bar = 100 μm. **(R)** Quantification of the percentage of pH3-positive and GFP-positive cells among GFP-positive cells in the cerebral cortex. Unpaired two-tailed t-test was used to compare different groups (N = 6 for each). Data are presented as mean±SEM. *P < 0.05, **P < 0.01, ***P < 0.001.

The proband in family_2 was a 5-year 6-month-old male with severe global developmental delay and congenital microcephaly. Born to non-consanguineous parents with no family medical history, the mother recalled a prenatal ultrasound indicating small fetal head circumference, though records were unavailable. He was born full-term via uncomplicated vaginal delivery. At 29 days old, he had a focal-to-secondary generalized epileptic seizure. Physical examination showed microcephaly (head circumference 31.5 cm, -3.4SD) and left thumb polydactyly, with normal muscle tone. At 1 year 1 month, EEG revealed multifocal epileptiform activity, mainly in bilateral occipital and temporal regions, including four right posterior temporal-originated seizures during sleep. Brain MRI showed cerebral dysgenesis, with pachygyria, corpus callosum thinning, and ventriculomegaly. Genital assessment noted right testis descent and left cryptorchidism. Valproic acid monotherapy failed to control his epilepsy. At 5 years 6 months, he remained severely developmentally delayed, unable to sit, crawl, or roll, and without meaningful speech. His head circumference was 45.7 cm (< -4.1SD), while height (110 cm) and weight (46.5 kg) were normal. All of the above phenotypic features can be found summarized in Table S1. Trio-WES identified a de novo heterozygous nonsense variant (NM_016359.4: c.1209C>G, p.Tyr403*) in *NUSAP1* exon 10.

These clinical findings, combined with the de novo variants in *NUSAP1*, suggest a potential pathogenic role of this gene in cortical development.

### NUSAP1 Deficiency Impairs NSPC Proliferation in the Developing Murine Cortex

Upon examining the expression profile of NUSAP1 in the UCSC databases, we observed that NUSAP1 was highly expressed in NSPCs (Fig. S1A). Additionally, IF analysis revealed that NUSAP1 was co-expressed with SOX2, a well-known marker of NSPCs, in hESC organoids and the E14.5 mouse brain (Fig. S1C, D).

To elucidate the functional role of NUSAP1 in neural development, we developed a *Nusap1* gene edited mouse model by targeting a specific segment within exon 10 of the *Nusap1* gene, corresponding precisely to the mutation site observed in patients with neurodevelopmental disorders, thus creating a clinically relevant system (Fig. 1F, G). Since both of the reported NUSAP1 mutations are located in the C-terminal domain, this genetically engineered mouse model is expected to produce C-terminal deficient NUSAP1 proteins, which plausibly replicate the C-terminal dysfunction of NUSAP1 observed in our patients. Throughout this study, these genetically engineered mice will be designated as *Nusap1*^+/Δ^ or *Nusap1*^Δ/Δ^, indicating heterozygosity or homozygosity for the targeted modification, respectively. First, to assess the molecular consequences of the genetic modification, we initially performed WB analysis to quantify NUSAP1 expression levels. The results showed that the expression of NUSAP1 protein in *Nusap1*^+/Δ^ and *Nusap1*^Δ/Δ^ mice was significantly lower than that in *Nusap1*^+/+^ mice (Fig. 1H, I). Furthermore, morphological analysis at embryonic day (E) 14.5 revealed significant reductions in both body size in *Nusap1*^+/Δ^ and *Nusap1*^Δ/Δ^ embryos compared to their *Nusap1*^+/+^ counterparts (Fig. 1J, K). Quantitative assessment of cerebral cortex size and cortical thickness revealed a significant reduction in both parameters in embryos of the two mutant genotypes compared to *Nusap1*^+/+^ embryos (Fig. 1L). Together, these findings demonstrate that our *Nusap1* gene-edited mouse model recapitulates key features of the human disorder and provides a relevant system for mechanistic investigation.

Building upon these observations, we next investigated the proliferation dynamics of NSPC populations in the developing cortices of *Nusap1*^+/Δ^ and *Nusap1*^Δ/Δ^ embryos. IF analysis revealed a significant reduction in the density of NUSAP1-positive cells, with mutant cortices displaying approximately 50% fewer positive cells compared to WT controls (Fig. 2A, B). To further characterize neural stem cells (NSCs) and NSPCs populations, we analyzed the expression of specific markers, demonstrating marked decreases in both SOX2-positive and PAX6-positive cell densities in mutant cortices relative to WT (Fig. 2C-F). Quantitative assessment of proliferative capacity through Ki67 immunostaining showed a substantial reduction in positive cells within mutant cortices compared to WT, indicating impaired mitotic activity (Fig. 2G, H). The number of mitotic cells, as indicated by phosphorylated histone H3 (pH3) positive cells, was significantly increased in *Nusap1*^+/Δ^ and *Nusap1*^Δ/Δ^ cortices compared to WT (Fig. 2I, J). To investigate the cellular mechanisms underlying these observations, we examined cell apoptosis using TUNEL assay. The results showed that apoptotic indices were elevated in mutant cortices, which was consistent with the observed cell cycle abnormalities (Fig. 2K, L). These abnormalities suggest that NUSAP1 play a crucial role in regulating the proliferation and expansion of NSPCs during brain development. The staining results of NUSAP1, Ki67, EdU, and TUNEL clearly demonstrated a significantly more marked impact in the homozygote group compared to the heterozygote group, strongly indicating the existence of a dose-dependent effect.

To specifically determine whether the observed functional abnormalities were directly caused by NUSAP1 deficiency rather than other potential factors, we performed in utero electroporation of NUSAP1 expression constructs into the developing brains of *Nusap1*^Δ/Δ^ embryos at E14.5 (Fig. 2M). Given the proximity of the mutations identified in our patients, we utilized the c.1213C>T mutation as a model system to investigate the functional consequences of mutations. Both WT and c.1213C>T mutant human *NUSAP1* mutations were cloned into the pCAGIG vector, which allows for simultaneous expression of the transgene and GFP reporter. This genetic rescue strategy allowed us to directly test the functional integrity of these constructs in restoring normal neural development, while providing definitive evidence that the observed defects were specifically mediated by NUSAP1 deficiency. Embryos were harvested three days post-electroporation, and brain tissues exhibiting GFP expression, indicating successful plasmid delivery and expression, were selected for subsequent analysis.

IF and WB analyses confirmed successful reintroduction of *NUSAP1* or c.1213C>T plasmid into *Nusap1*^Δ/Δ^ embryos at E17.5, as evidenced by colocalization of NUSAP1-positive and GFP-positive cells. Importantly, the electroporation efficiency was comparable between the two constructs (Fig. 2N). Notably, quantitative analysis revealed that the p.Gln405* NUSAP1 exhibited higher expression levels of the truncated protein compared to WT NUSAP1 in the embryonic cerebral cortex (Fig. 2P, Q). Despite this increased expression, functional assessment demonstrated that the p.Gln405* variant was significantly impaired in its ability to rescue NUSAP1 deficiency. PH3 immunostaining revealed that while WT NUSAP1 effectively reduced the mitotic cell population, indicating successful functional recovery, the p.Gln405* variant failed to normalize the abnormal mitotic index (Fig. 2O). Quantitative analysis of pH3-positive/GFP-positive cells among GFP-positive populations further confirmed the reduced functional capacity of the truncated protein (Fig. 2R). Collectively, these findings demonstrate that the p.Gln405* mutation disrupts cortical development in mouse embryos through dysregulation of mitotic processes, highlighting the essential role of intact NUSAP1 function in normal brain development.

### NUSAP1 Mutations Induce Spindle Dynamics Aberrations, Cell Cycle Disruption, and Apoptosis

Although our study has established a connection between NUSAP1 and microcephaly, the molecular mechanisms underlying its function remain poorly understood. To investigate the molecular role of NUSAP1 in disease pathogenesis, we generated HA-tagged plasmids containing either WT *NUSAP1* or various *NUSAP1* mutations for overexpression studies in cultured cells. Additionally, to facilitate cellular localization analysis, we constructed GFP-tagged NUSAP1 plasmids for ICC. Given previous reports implicating NUSAP1 in spindle assembly(22), we transfected HEK293T cells with GFP-tagged NUSAP1 and performed co-staining with α-tubulin to visualize spindle morphology and γ-tubulin to mark the centrosome in metaphase cells. Notably, *NUSAP1* mutations expression resulted in severe spindle abnormalities, including significantly shortened spindle lengths and the formation of multipolar spindles within individual cells (Fig. 3A-D). Chromosome alignment was also disrupted in mutant cells, with chromosomes failing to form the characteristic metaphase plate observed in control cells (Fig. 3A). RT-qPCR and WB analyses of HEK293T cells transfected with HA-tagged NUSAP1 demonstrated that *NUSAP1* mutations showed significantly higher expression levels at both the transcriptional and translational levels compared to the control (Fig. 3E-G). Actinomycin D treatment experiments demonstrated that this increased expression was attributable to enhanced mRNA stability of *NUSAP1* mutations (Fig. 3H). This not only explained the elevated mutant protein expression in vitro, but also implies that other mechanisms may account for the reduced NUSAP1 expression in in vivo-edited mice.

**Figure 3.**
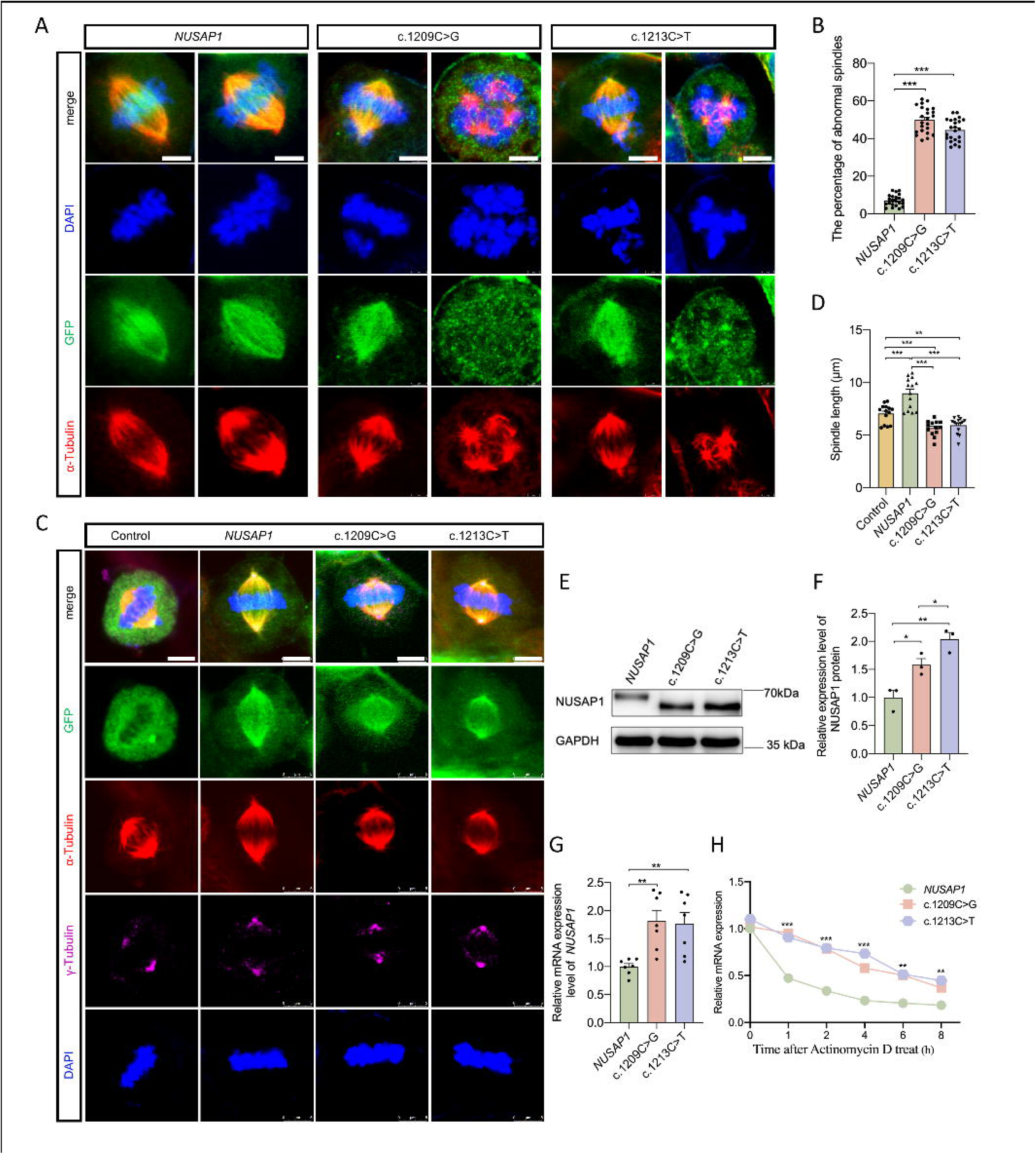
NUSAP1 mutations alter spindle structure, protein and mRNA expression in HEK293T cells. **(A)** Immunofluorescent images of HEK293T cells: nuclei (DAPI-blue), GFP (green), α-Tubulin (red), merged panels were shown above. Scale bar = 5LJμm. **(B)** Quantification of the percentage of cells with abnormal spindles. **(C)** Immunofluorescent images of HEK293T cells: GFP (green), α-Tubulin (red), γ-Tubulin (purple) and nuclei (DAPI-blue), merged panels were shown above. Scale bar = 5LJμm. **(D)** Quantification of the spindle length in each group. **(E)** Western blot analysis was performed to detect the NUSAP1 expression in HEK293T cells transfected with *NUSAP1*, c.1209C>G or c.1213C>T plasmids. **(F)** Quantification of the NUSAP1 protein expression levels in three groups. **(G)** Quantification of the relative mRNA expression level in three groups. **(H)** Quantification of relative mRNA expression levels in HEK293T cells in response to treatment with 10 μg/ml Actinomycin D. Unpaired two-tailed t-test compared different experimental groups for significant differences. Data are shown as mean±SEM. \**P*<0.05, \*\**P*<0.01, \*\*\**P*<0.001.

To further investigate the impact of NUSAP1 overexpression on cell cycle progression, we performed immunofluorescence staining for pH3, a mitotic marker, and 5-ethynyl-2’-deoxyuridine (EdU), a marker of DNA synthesis, in HEK293T cells transfected with either control, WT *NUSAP1* or *NUSAP1* mutations. Quantification of pH3-positive cells revealed no significant difference between mock transfection control cells and those transfected with *NUSAP1* (Fig. 4A-B). However, cells transfected with the *NUSAP1* mutations exhibited a significant increase in the proportion of pH3-positive cells, indicating an elevated mitotic index (Fig. 4A-B). Conversely, the percentage of EdU-positive cells remained consistent across all experimental groups, suggesting that NUSAP1 overexpression does not directly perturb cell proliferation (Fig. 4A, C). Notably, the ratio of pH3-positive to EdU-positive cells was significantly higher in cells transfected with *NUSAP1* mutations compared to those transfected with *NUSAP1* (Fig. 4D). Given that pH3 specifically marks the mitotic phase of the cell cycle, while EdU labels the S phase, this elevated ratio suggests a potential disruption of cell cycle checkpoint regulation. Consequently, we hypothesized that NUSAP1 mutations might induce spindle assembly checkpoint (SAC). To directly test whether SAC is activated, we measured mRNA levels of the checkpoint regulators BUBR1 and MAD2. BUBR1, along with MAD2, is a key protein involved in the SAC. The RT-qPCR results demonstrated a significant increase in the expression of both *BUBR1* and *MAD2* in cells transfected with the *NUSAP1* mutations, supporting our hypothesis (Fig. 4E, S2). Next, cell cycle distribution was analyzed by flow cytometry in HEK293T cells transfected with control, *NUSAP1*, c.1209C>G, or c.1213C>T plasmids. Notably, the proportion of cells in S phase was decreased in cells transfected with the *NUSAP1* mutations compared to control. Conversely, the proportion of cells in the G2/M phase was significantly increased in cells transfected with the *NUSAP1* mutations (Fig. 4F-G). Collectively, these results suggested that the *NUSAP1* mutations-transfected cells displayed a potential arrest in the mitotic phase of the cell cycle. Furthermore, we assessed apoptosis by flow cytometry in HEK293T cells transfected with the same plasmids, and found that transfection with both mutant plasmids resulted in a significant increase in cell apoptosis (Fig. 4H, I). These findings demonstrate that *NUSAP1* mutations disrupt mitotic progression by activating the spindle assembly checkpoint, leading to cell cycle arrest and apoptosis.

**Figure 4.**
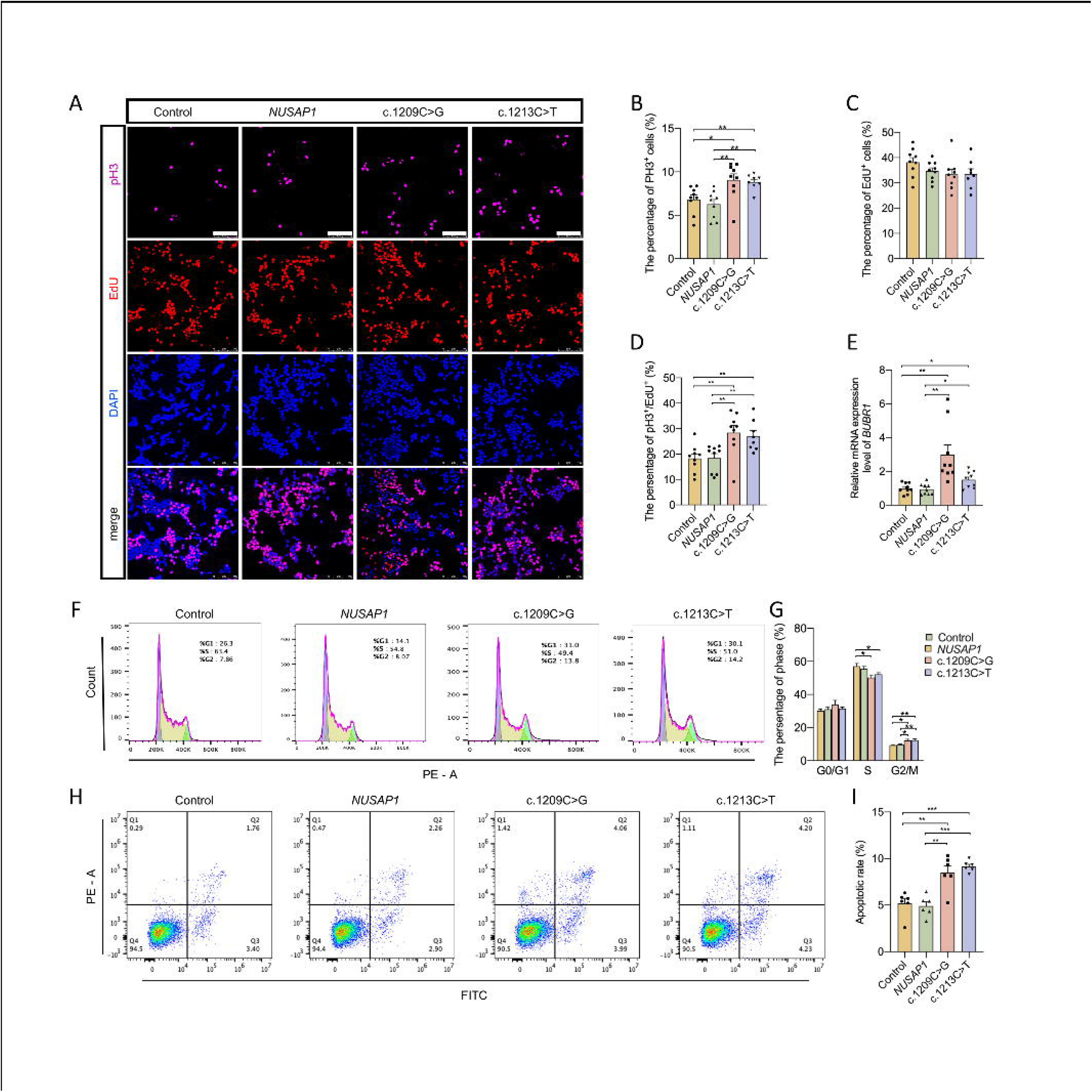
Effect of NUSAP1 and its mutations on mitotic activity, cell cycle distribution, and apoptosis in HEK293T Cells. **(A)** Representative images of pH3 and EdU staining in HEK293T cells overexpressing NUSAP1. The merged images were shown below. Scale bar = 100LJμm. **(B-D)** Quantification of the percentages of pH3-positive cells, EdU-positive cells, and cells with pH3/EdU signals, respectively. **(E)** The relative mRNA expressions of *BUBR1* was calculated by comparing their expression values with those in control cells, which were normalized to 1. **(F, G)** Cell cycle was analyzed by flow cytometry in HEK293T cells transfected with control, *NUSAP1*, c.1209C>G or c.1213C>T plasmids, showing the distribution of cells in G0/G1, S and G2/M phases. **(H, I)** Apoptosis was analyzed by flow cytometry in HEK293T cells transfected with control, *NUSAP1*, c.1209C>G or c.1213C>T plasmid. Unpaired two-tailed t-test was used to compare different experimental groups for significant differences. Data are shown as mean±SEM. \**P* < 0.05, \*\**P* < 0.01, \*\*\**P* < 0.001.

### Disease-Associated NUSAP1 Mutations Fail to Rescue KIF2C-Induced Mitotic Defects

Given that our *NUSAP1* mutations are unable to fulfill this fundamental cellular function, we sought to investigate the molecular mechanisms connecting NUSAP1 to spindle and chromosome dynamics during mitosis. More intriguingly, upon examining the expression profile of NUSAP1 in mouse and human brain databases (Fig. S1A, B), we noted that KIF2C exhibits a strong co-expression pattern with NUSAP1, particularly in NSPCs, which prompted our focus on KIF2C. Previous studies have identified KIF2A as a spindle-associated protein(28). KIF2C, a member of the same subfamily, shares structural similarity but its precise function and potential involvement in microcephaly remain unclear. Therefore, we aimed to elucidate the functional relationship between these two proteins.

Initially, we examined immunofluorescent images of the E14.5 mouse embryo cerebral cortex stained for NUSAP1 and KIF2C to determine their co-expression and localization. This analysis revealed clear colocalization of NUSAP1 and KIF2C in the ventricular zone (VZ) and subventricular zone (SVZ) region of the E14.5 mice cortex (Fig. 5A). To investigate their molecular interaction, we co-transfected HEK293T cells with a GFP-tagged KIF2C plasmid and an HA-tagged NUSAP1 plasmid. ICC analysis of these transfected cells demonstrated colocalization of KIF2C and NUSAP1 within the nucleus and, notably, at the spindle apparatus, which was identified by α-tubulin staining (Fig. 5B). To determine whether KIF2C and NUSAP1 interact, we performed reciprocal Co-IP assays in co-transfected HEK293T cells. Both approaches demonstrated that KIF2C and NUSAP1 interact (Fig. 5C, D). Subsequently, Co-IP assays with KIF2C and NUSAP1 mutations revealed that both mutations disrupted this interaction, preventing them from interacting with KIF2C to exert its function (Fig. 5E).

**Figure 5.**
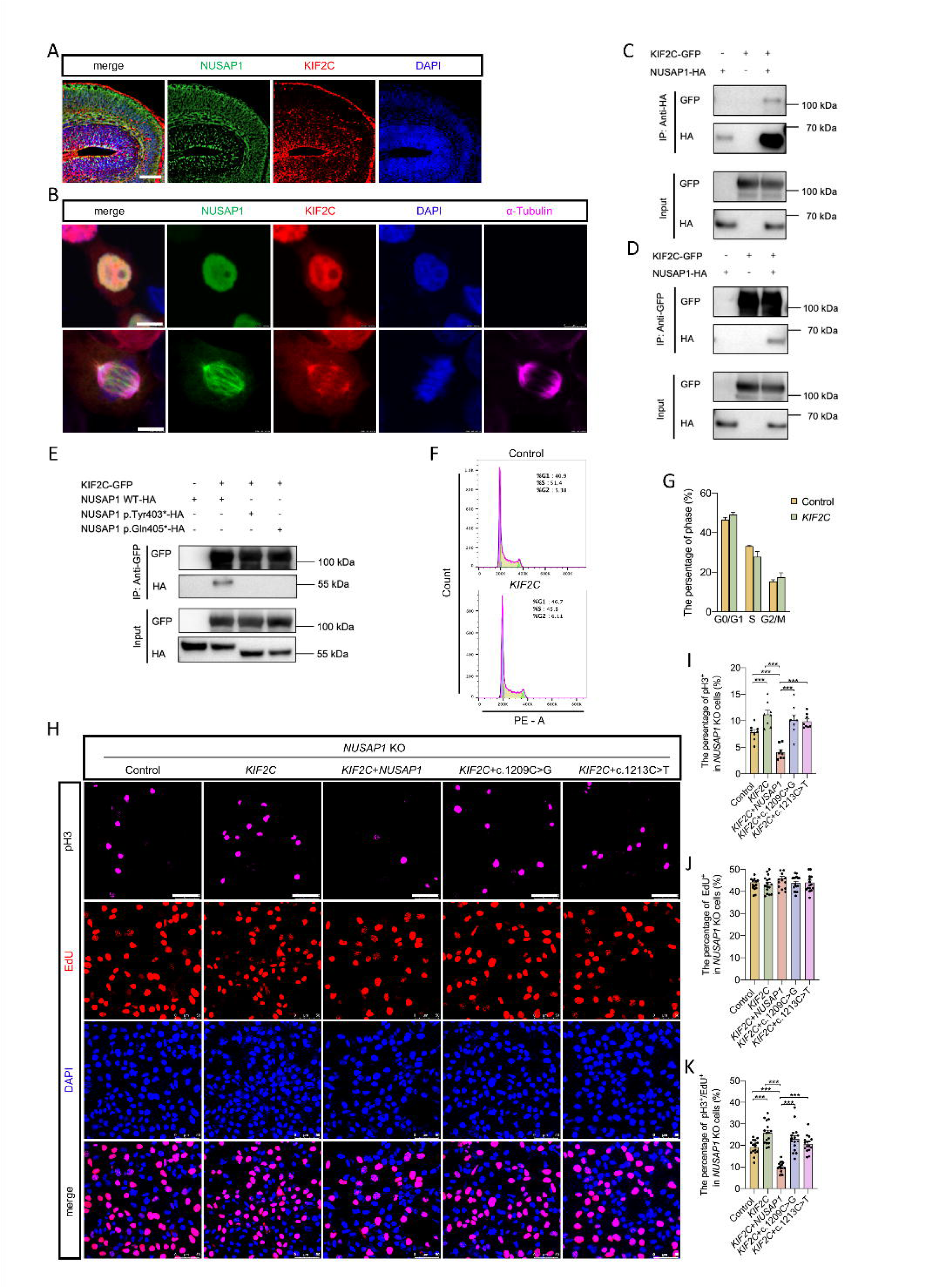
Interaction, co-localization and functional impact of NUSAP1 and KIF2C in cerebral cortex and HEK293T Cells. **(A)** Immunofluorescent images of the cerebral cortex stained for NUSAP1 (green) and KIF2C (red) were used to detect the co-expression and co-localization of these two proteins. DAPI was used to stain the nuclei (blue). Merged images of the NUSAP1, KIF2C, and DAPI channels are shown on the left. Scale bar = 100 μm. **(B)** Immunofluorescent images of HEK293T cells, with NUSAP1 stained green and KIF2C stained red, were obtained to detect the co-expression and co - localization of these two proteins. DAPI, in blue, was utilized to stain the nuclei. Additionally, α-Tubulin was stained purple to mark the spindles. Scale bar = 10 μm. **(C, D)** HEK293T cells were transfected with *KIF2C*-GFP and *NUSAP1*-HA plasmids, and then both forward and reverse Co-IP assays were used to detect the interaction between NUSAP1 and KIF2C. **(E)** HEK293T cells were transfected with plasmids encoding *KIF2C*-GFP, *NUSAP1*-HA, or two NUSAP1 mutants. Subsequently, Co-IP assays were performed to detect the interactions between KIF2C and NUSAP1, as well as those between KIF2C and the two NUSAP1 mutant proteins. **(F, G)** Cell cycle was analyzed by flow cytometry in HEK293T cells transfected with control or *KIF2C* plasmids, showing the distribution of cells in G0/G1, S and G2/M phases. **(H)** Representative images of pH3 and EdU staining in *NUSAP1* KO HEK293T cells overexpressing KIF2C and NUSAP1. The merged images were shown below. Scale bar = 50LJμm. **(I-K)** Quantification of the percentages of pH3-positive cells, EdU-positive cells, and cells with pH3/EdU signals, respectively. Unpaired two-tailed t-test was used to compare different experimental groups for significant differences. Data are shown as mean±SEM. \**P* < 0.05, \*\**P* < 0.01, \*\*\**P* < 0.001.

Previous research has demonstrated that KIF2C interacts directly with the kinetochore to regulate chromosome alignment during mitosis(29). Given our observation that KIF2C interacts with NUSAP1, and their similar subcellular localization in embryonic cortex, we sought to determine whether KIF2C plays a role in cell cycle progression and mitosis similar to NUSAP1, and to elucidate their regulatory relationship. Cell cycle analysis of HEK293T cells transfected with control or *KIF2C* plasmids revealed no significant differences in cell cycle phase distribution (Fig. 5F, G). We hypothesized that the effect of KIF2C overexpression could be attenuated by endogenous NUSAP1. This is because endogenous NUSAP1 might interact with the transfected GFP-KIF2C and modulate its function. To exclude the influence of endogenous *NUSAP1* expression, we generated a *NUSAP1* KO HEK293T cell line using CRISPR-Cas9 gene editing. Knockout efficiency was confirmed by Western blotting (Fig. S2A). Cell cycle distribution analysis by flow cytometry revealed that *NUSAP1* KO also resulted in cell cycle dysregulation, mirroring the effects observed in HEK293T cells transfected with NUSAP1 mutations (Fig. S2B, C). And the cell cycle distribution was also analyzed in HEK293T cells transfected with control, *NUSAP1*, c.1209C>G, or c.1213C>T plasmids. Notably, the proportion of cells in G2/M phase was decreased in *NUSAP1* KO cells transfected with the *NUSAP1* mutations compared to *NUSAP1* KO controls. Conversely, the mutations had no significant rescue effect compared to the control (Fig. S2D, E). For apoptosis analysis, only cells transfected with WT *NUSAP1* exhibited a reduction in apoptosis to normal levels, whereas the mutations had no significant effect compared to the control (Fig. S2F, G). Therefore, to investigate the regulatory relationship between these two proteins, we performed pH3 and EdU staining in *NUSAP1* KO HEK293T cells overexpressing *KIF2C* and *NUSAP1* (Fig. 5H). Interestingly, *NUSAP1* KO cells transfected with *KIF2C* showed a significant increase in the proportion of pH3-positive cells (Fig. 5I). To compare different cell populations and exclude potential confounding factors, we normalized pH3-positive cell ratios to EdU-positive cell ratios, which showed no difference across cell lines (Fig 5J). Notably, this increase in pH3-positive/Edu-positive cells in *NUSAP1* KO cells could be rescued by co-transfection with WT *NUSAP1*, but not by *NUSAP1* mutations (Fig. 5K).

Consistent with these findings, cell cycle analysis revealed that *NUSAP1* KO HEK293T cells transfected with *KIF2C*, *KIF2C*+c.1209C>G, and *KIF2C*+c.1213C>T plasmids showed a decrease in G0/G1 phase and increase in G2/M phase, whereas those transfected with control and *KIF2C*+*NUSAP1* exhibited the opposite effect (Fig. 6A, B). As described earlier, we propose that NUSAP1 dysfunction may promote SAC, leading to abnormal spindle formation. In cells undergoing SAC, BUBR1, a key mitotic checkpoint protein, localizes to the vicinity of misaligned chromosomes. This localization is characterized by BUBR1-positive cells in ICC assays. To further investigate the influence of *NUSAP1* and its mutations on SAC, we performed ICC on *NUSAP1* KO HEK293T cells transfected with control, *KIF2C*, *KIF2C*+*NUSAP1*, *KIF2C*+c.1209C>G, or *KIF2C*+c.1213C>T plasmids (Fig. 6C). Our results demonstrate that overexpression of *KIF2C* in *NUSAP1* KO HEK293T cells significantly increased the percentage of BUBR1-positive cells (Fig. 6D-F), indicating an elevated level of SAC. Importantly, this abnormal SAC activation was rescued by co-transfection with *NUSAP1*, while the *NUSAP1* mutations failed to reverse this effect (Fig. 6D-F). Results of apoptosis analysis showed that *KIF2C* overexpression also induced a significant increase in the apoptotic rate in *NUSAP1* KO HEK293T cells. This elevated apoptosis was rescued by co-transfection with *NUSAP1*, whereas the NUSAP1 mutations failed to abrogate the increase (Fig. 6G, H). These findings suggest that functional NUSAP1 is required to suppress KIF2C-induced SAC and apoptosis.

**Figure 6.**
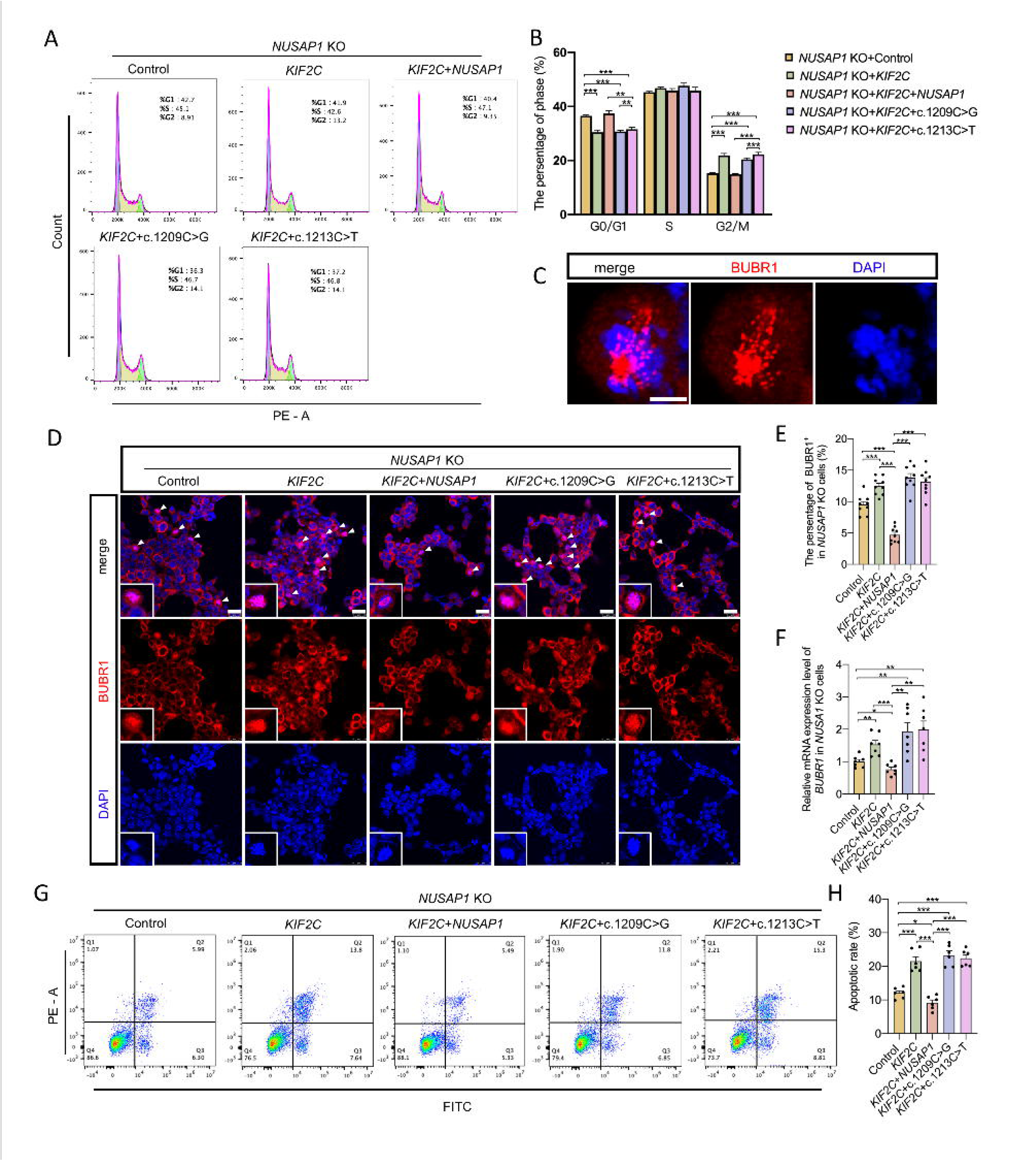
Impact of KIF2C and NUSAP1 mutations on cell cycle progression, spindle assembly checkpoint and apoptosis in *NUSAP1* KO HEK293T Cells. **(A, B)** Cell cycle was analyzed by flow cytometry in *NUSAP1* KO HEK293T cells transfected with control, *KIF2C*, *KIF2C*+*NUSAP1*, *KIF2C*+c.1209C>G or *KIF2C*+c.1213C>T plasmids, showing the distribution of cells in G0/G1, S and G2/M phases. **(C-E)** Immunofluorescent images of the *NUSAP1* KO HEK293T cells transfected with control, *KIF2C*, *KIF2C*+*NUSAP1*, *KIF2C*+c.1209C>G or *KIF2C*+c.1213C>T plasmids and stained for BUBR1 were used to analyze the function of the spindle assembly checkpoint. DAPI was used to stain the nuclei (blue). Merged images of the BUBR1 and DAPI channels are shown above. The white triangles indicate BUBR1-positive cells. Scale bar = 25 μm. **(F)** The relative mRNA expression of *BUBR1* was calculated by comparing their expression values with those in control cells, which were normalized to 1. **(G, H)** Apoptosis was analyzed by flow cytometry in *NUSAP1* KO HEK293T cells transfected with control, *KIF2C*, *KIF2C*+*NUSAP1*, *KIF2C*+c.1209C>G or *KIF2C*+c.1213C>T. Unpaired two tailed t-test was used to compare different experimental groups for significant differences. Data are shown as mean±SEM. \**P* < 0.05, \*\**P* < 0.01, \*\*\**P* < 0.001.

Together, these findings reveal that KIF2C acts downstream of NUSAP1 in a signaling pathway. Pathogenic *NUSAP1* mutations disrupt this regulatory axis, leading to abnormal cell proliferation and mitosis—phenotypes that likely contribute to the pathogenesis of microcephaly.

### The AURKA-NUSAP1 Signaling Axis Regulates KIF2C Function and Cell Cycle Progression

Previous research has established that phosphorylation of NUSAP1 is crucial for its proper function in cell cycle progression. Among the kinases implicated in this process, Aurora kinase A (AURKA) has been shown to play a significant role in regulating mitosis(30, 31). We first examined the expression and localization of AURKA in embryonic cortex and found clear colocalization of NUSAP1 and AURKA, confirming their connection (Fig. 7A). Interestingly, we found that in *Nusap1*^+/Δ^ and *Nusap1*^Δ/Δ^ cortices, AURKA-positive cells ratio was also decreased (Fig. 7B, C). To further investigate the function of AURKA in NUSAP1 regulation, we generated a Flag-tagged AURKA plasmid. Next, HEK293T cells were transfected with AURKA-Flag and NUSAP1-HA plasmids, and reciprocal Co-IP assays were performed to detect the interaction between NUSAP1 and AURKA. Co-IP results showed positive results in both experiments, confirming an interaction between NUSAP1 and AURKA (Fig. 7D, E). Then, we performed Co-IP in HEK293T cells transfected with plasmids encoding AURKA-Flag and the two NUSAP1 mutations. However, results showed that the mutations did not disrupt the interaction between AURKA and NUSAP1 (Fig. 7F). Moreover, to investigate whether the interaction persists but the NUSAP1 mutations are not correctly modified by AURKA, we used MLN8237 to inhibit AURKA kinase activity and then detect NUSAP1 downstream interaction as described previously. Surprisingly, we found that MLN8237 significantly decreased the NUSAP1-KIF2C interaction in co-transfected HEK293T cells (Fig. 7G).

**Figure 7.**
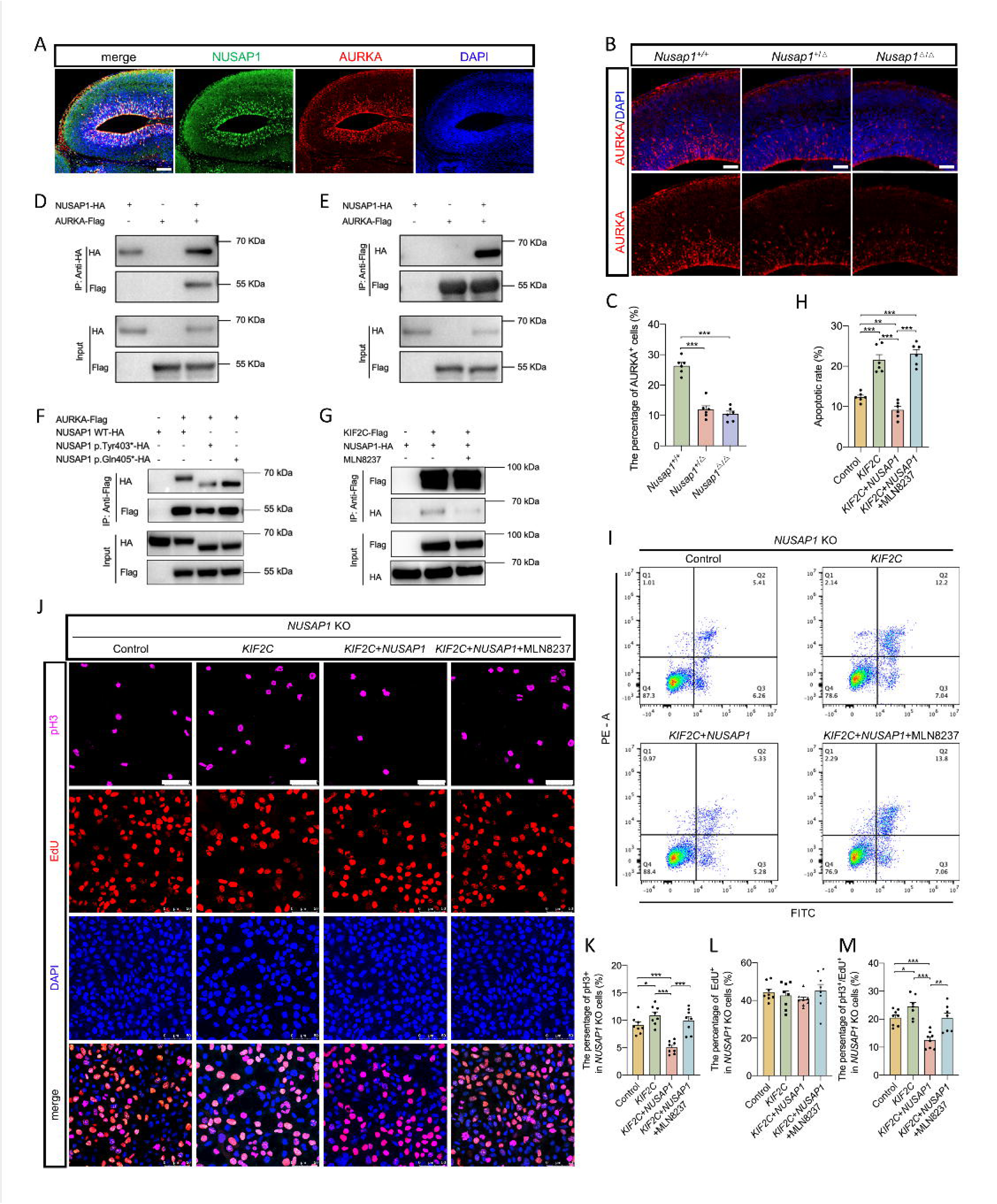
AURKA regulates the interaction and function of NUSAP1-KIF2C. **(A)** Immunofluorescent images of the cerebral cortex stained for NUSAP1 (green) and AURKA (red) were used to detect the co-expression and co-localization of these two proteins. DAPI was used to stain the nuclei (blue). Merged images of the NUSAP1, AURKA, and DAPI channels are shown on the left. Scale bar = 100 μm. **(B)** Immunofluorescent images of the cerebral cortex stained for AURKA (red) to detect AURKA-positive cells. The merged images of the AURKA and DAPI channels were shown above. Scale bar = 50 μm. **(C)** Quantification of the percentage of AURKA-positive cells in the cerebral cortex. **(D, E)** HEK293T cells were transfected with AURKA-Flag and NUSAP1-HA plasmids, and then both forward and reverse Co-IP assays were used to detect the interaction between NUSAP1 and AURKA. (**F)** HEK293T cells were transfected with plasmids encoding AURKA-Flag, NUSAP1-HA, or two NUSAP1 mutant proteins. Subsequently, Co-IP assays were performed to detect the interactions between AURKA and NUSAP1, as well as those between AURKA and the two mutant proteins. **(G)** HEK293T cells were transfected with plasmids encoding KIF2C-Flag and NUSAP1-HA. Subsequently, MLN8237 was added to inhibit the Aurora kinase. Co-IP assays were performed to detect the interactions between KIF2C and NUSAP1, both in the absence and presence of MLN8237. **(H, I)** Apoptosis was analyzed by flow cytometry in *NUSAP1* KO HEK293T cells transfected plasmids encoding control, *KIF2C*, *KIF2C* + *NUSAP1*. The percentages of apoptotic cells were quantified both in the absence and presence of MLN8237. **(J-M)** Representative images of pH3 and EdU staining in *NUSAP1* KO HEK293T cells which were transfected with control, *KIF2C*, *KIF2C*+*NUSAP1* and *KIF2C*+*NUSAP1*+MLN8237. The merged images were shown below. Scale bar = 50LJμm. Quantification of the percentages of pH3-positive cells, EdU-positive cells, and cells with pH3/EdU signals, respectively. Unpaired two-tailed t-test was used to compare different experimental groups for significant differences. Data are shown as mean±SEM. \**P* < 0.05, \*\**P* < 0.01, \*\*\**P* < 0.001.

Then, apoptosis was analyzed in *NUSAP1* KO HEK293T cells transfected with plasmids encoding control, *KIF2C*, and *KIF2C*+*NUSAP1*, as previously described. Apoptosis analysis revealed that the addition of MLN8237 prevented *NUSAP1* from rescuing the increase in apoptosis caused by *KIF2C* overexpression (Fig. 7H, I). Moreover, we also performed pH3 and EdU staining in *NUSAP1* KO HEK293T cells transfected with control, *KIF2C*, *KIF2C*+*NUSAP1*, and treated *KIF2C*+*NUSAP1* group with MLN8237. Consistent with the apoptosis data, MLN8237 treatment abolished the ability of *NUSAP1* to rescue the effects of overexpressed *KIF2C* in *NUSAP1* KO HEK293T cells (Fig. 7J-M).

These findings define an AURKA–NUSAP1–KIF2C signaling axis essential for mitotic progression. Disruption of this phosphorylation-dependent pathway—whether by kinase inhibition or disease-causing mutations—leads to mitotic arrest and apoptosis, providing mechanistic insight into the pathogenesis of microcephaly.

## Discussion

Microcephaly, a severe neurological disorder, represents a significant medical challenge, as the molecular mechanisms underlying its pathogenesis remain incompletely understood. In this study, we report the identification of two de novo mutations in NUSAP1—c.1209C>G (p.Tyr403*) and c.1213C>T (p.Gln405*)—in patients presenting with microcephaly. Notably, during our global patient-matching efforts via GeneMatcher, we identified that *Mo, A. et al.* had independently reported the same locus truncating mutation, c.1209C>A (p.Tyr403*), in two unrelated families affected by microcephaly. However, a formal collaboration could not be established due to the extended timeline required for the generation of animal models. Consequently, their findings were published as case reports in Clinical Genetics in July 2023(32). Given the potential role of *NUSAP1* in critical neurodevelopmental processes, including neural crest cell migration and morphogenesis(32–34), and the identification of *NUSAP1* mutations in microcephaly patients, we investigated the role of NUSAP1 in brain development and the microcephaly disease mechanism. Using *Nusap1*^+/Δ^ and *Nusap1*^Δ/Δ^ mouse models, and *NUSAP1* KO HEK293T cells, we examined the consequences of *NUSAP1* dysfunction. Our findings from ICC and cell cycle analyses indicate that NUSAP1 plays a critical role in spindle dynamics and cell mitosis. Disruption of NUSAP1 function, such as that caused by the identified mutations, impairs its interaction with KIF2C, leading to aberrant microtubule flux and abnormal spindle assembly. Furthermore, we confirmed the necessity of AURKA-mediated dynamic regulation of NUSAP1 in the NUSAP1-KIF2C pathway. Our results suggest that NUSAP1 mutations disrupt spindle assembly by impairing KIF2C function through a mechanism involving aberrant AURKA regulation. This leads to cell cycle and mitosis disturbances, resulting in decreased NSPCs numbers during development, exhaustion of the stem cell pool, and ultimately, microcephaly.

WB analysis revealed an increase in *NUSAP1* variant protein expression in the cerebral cortex of *Nusap1*^Δ/Δ^ mice (Fig. 2P, Q) and in vitro HEK293T cells (Fig. 3E, F). Based on this finding, we hypothesized that this increase was due to increased mRNA stability caused by the *NUSAP1* mutations. To further validate this hypothesis, we examined the mRNA degradation rates of HEK293T cells transfected with different *NUSAP1* plasmids. As shown in Fig. 3H, these cells exhibited varying mRNA degradation rates, suggesting that the mutations indeed alter NUSAP1 mRNA stability. Moreover, although cell apoptosis analysis revealed an increased apoptosis rate in HEK293T cells transfected with different *NUSAP1* variant plasmids, the total cell number might not be significantly affected in the short term. In the mouse model, WB analysis revealed that while gene editing did not result in premature termination of protein translation, the protein levels translated from the sequence with a 13-bp deletion were significantly lower than those of the WT (Fig. 1H, I). This finding contrasts with the upregulated expression of truncated mutant proteins described above. We propose that, in addition to potential differences in mRNA degradation levels, the more critical factor is the abnormal mitosis and cell proliferation caused by alterations in the C-terminal sequence of NUSAP1. These disruptions result in a decline in the population of proliferative NSPCs, a conclusion buttressed by our observation of reduced numbers of NUSAP1-positive cells, as well as SOX2-positive and PAX6-positive cells, in the mouse cortex (Fig. 2A - F). Since *NUSAP1* is predominantly expressed in dividing cells, these cellular alterations ultimately exert a profound impact on its overall protein expression levels(34–36). This finding provides an underlying explanation for the discrepancy observed between the in vivo Ki67 staining, which indicated changes in cell proliferation (Fig. 2G, H), and the in vitro EdU staining, where no significant differences were detected (Fig. 4A, C). It is plausible that within the 48-hour period, while the cells were prompted to enter the mitotic phase, they failed to fully initiate or complete substantial DNA replication.

AURKA, a serine/threonine kinase, plays a critical role in regulating several mitotic events, including mitotic entry, spindle assembly, and chromosome segregation(37). Its expression and activity are typically low during interphase and peak during mitosis. Given that NUSAP1 mutations lead to less successful mitosis, the number of cells capable of proliferation would decrease during cortical development. Consistent with this, we observed a reduction in AURKA-positive cells in the cortices of *Nusap1*^+/Δ^ and *Nusap1*^Δ/Δ^ mice (Fig. 7B, C). This would result in a reduction in the total number of proliferative cells, ultimately leading to the observed decrease in AURKA-positive cells in the cortex. Notably, AURKA has been predominantly studied in the context of cancer biology(38–40), with limited research exploring its role in neural development and neurological disorders. Previous research has demonstrated that AURKA-mediated phosphorylation of NUSAP1 is essential for maintaining its interaction with KIF family proteins, thereby ensuring proper spindle microtubule flux(20). This regulatory mechanism represents a dynamic equilibrium between AURKA and NUSAP1. Our findings establish a direct mechanistic link between the interaction of AURKA and NUSAP1 in cell cycle regulation and the pathogenesis of microcephaly, demonstrating their coordinated role in the regulation of mitotic spindle assembly in NSPCs.

Beyond AURKA, our data show that KIF2C, a key regulator of microtubule flux and spindle length control(41), is a critical downstream effector of NUSAP1. KIF2C, also known as MCAK, depolymerizes kinetochore microtubules and is negatively regulated by NUSAP1(25). During metaphase, the interaction between NUSAP1 and KIF2C reduces KIF2C depolymerization activity and promotes kinetochore microtubule stability(25). Consistent with this, our results showed that *KIF2C* overexpression in *NUSAP1* KO HEK293T cells led to abnormal cell cycle progression and mitosis, effects that were rescued by co-transfection with WT *NUSAP1*, but not by *NUSAP1* variants, suggesting a disruption of NUSAP1-KIF2C regulation by these variants. Furthermore, we observed that disruption of AURKA-mediated regulation of NUSAP1 also disrupted NUSAP1-KIF2C regulation, leading to abnormal cell proliferation and mitosis. Collectively, these results indicate that the AURKA-NUSAP1-KIF2C regulatory pathway is crucial for proper spindle formation and microtubule function, thereby ensuring successful cell mitosis. Within this pathway, NUSAP1 acts as a key regulator, linking different regulatory components. In this study, we established a novel connection between AURKA, NUSAP1, and KIF2C, elucidating their functional interplay during brain development.

We also observed that *KIF2C* overexpression in HEK293T cells did not directly induce cell cycle abnormalities similar to those seen with *KIF2C* overexpression in *NUSAP1* KO cells. Given our finding that KIF2C function is regulated by NUSAP1, we hypothesize that endogenous NUSAP1 in normal HEK293T cells is sufficient to regulate the overexpressed KIF2C. Specifically, the interaction between NUSAP1 and KIF2C, which deactivates KIF2C, may neutralize the effect of the excess KIF2C. However, overexpression of NUSAP1 variants in normal HEK293T cells did induce significant abnormalities in cell mitosis and apoptosis. We propose that, unlike the *KIF2C* overexpression model where endogenous NUSAP1 can compensate, overexpressed NUSAP1 variants disrupt the function of endogenous NUSAP1 in two ways. First, NUSAP1 variants may competitively inhibit AURKA binding, thus preventing proper NUSAP1 phosphorylation, which is essential for the subsequent NUSAP1-KIF2C interaction. The resulting insufficient phosphorylated NUSAP1 would then fail to adequately deactivate KIF2C, leading to abnormalities. Second, NUSAP1 variants, while still able to localize to spindle microtubules, may compete with endogenous NUSAP1 for critical binding sites (such as those on minus-end microtubules where NUSAP1 interacts with KIF2C). This competition could disable the ability of endogenous NUSAP1 to perform its essential role. Altogether, these different responses to KIF2C and NUSAP1 overexpression further support our conclusion that spindle regulation occurs via an AURKA-NUSAP1-KIF2C pathway. However, these hypotheses warrant further investigation, and additional studies are required to elucidate the precise role of KIF2C in microcephaly. Furthermore, given that both NUSAP1 and KIF2C are specifically expressed in NSPCs during brain development, we hypothesize that these two critical factors may play a more extensive co-regulatory role in cortical formation, which also merits further exploration.

Our study highlights the cell cycle disruption observed in NUSAP1 dysfunction models, including the *Nusap1* gene edited mouse model and NUSAP1-deficient HEK293T cells. In these models, we also found altered pH3-positive cell ratios and BUBR1 expression, indicating a damaged cell cycle. Since pH3 marking cells entering mitosis, specifically at the G2/M phase(42), and BUBR1 as a key component of SAC, these findings suggest that NUSAP1 dysfunction activates the SAC, potentially leading to increased apoptosis. While NUSAP1 may not directly cause this phenomenon, our data indicate that it functions as a crucial regulator within the AURKA-NUSAP1-KIF2C pathway. Disruptions in this pathway impair normal microtubule flux during mitosis, which could in turn activate the SAC(43). This activation likely contributes to the observed decrease in NSPCs numbers, ultimately leading to premature depletion of the NSPC pool and the microcephaly phenotype. Inspired by these findings, we speculate that SAC inhibition might offer a potential therapeutic strategy for microcephaly caused by NUSAP1 dysfunction. However, given the critical role of SAC in all cell types, careful dosage optimization would be necessary to avoid potential drug toxicity(44). The observation that WT NUSAP1 overexpression rescued the abnormal phenotype in our mouse model suggests that direct administration of WT NUSAP1 may be a more viable clinical approach. Furthermore, the finding that mutant NUSAP1 disrupts its interaction with KIF2C suggests that a molecule mimicking WT NUSAP1 to promote this interaction could also be a therapeutic strategy.

Several limitations in our study warrant mention. First, our analysis included only two de novo NUSAP1 mutants from two patients, which is insufficient for drawing broad conclusions about the entire human population. Larger population studies are needed to fully elucidate the detailed mechanisms of *NUSAP1* in microcephaly. Additionally, several questions remain unaddressed in our research. We did not investigate the pathways influenced by *NUSAP1* variants or NUSAP1 dysfunction. Given the importance of cell proliferation and apoptosis in development, NUSAP1 dysregulation may disrupt not only KIF2C but also other downstream pathways, such as cell migration(45, 46). Future research employing metabolomics or fluxomics could explore this. Although we found that *NUSAP1* mutations lead to abnormal spindle formation and cell mitosis, technical limitations prevented us from directly observing dynamic changes in microtubules. Finally, we observed that our *Nusap1*-edited mouse model exhibited reproductive abnormalities, which was unexpected. Because reproduction requires cell cycle progression to produce functional germline cells(47), NUSAP1 dysfunction might disrupt their formation. Further research is needed to confirm whether this phenotype results from NUSAP1-mediated germline-specific dysfunction.

In conclusion, our study provides compelling evidence that NUSAP1 represents a previously unrecognized genetic determinant in microcephaly. Through comprehensive in vitro and in vivo analyses, we have demonstrated that de novo mutations in NUSAP1 disrupt the finely-tuned AURKA-NUSAP1-KIF2C regulatory pathway. This disruption initiates a cascade of cellular events, including mitotic abnormalities and premature exhaustion of NSPCs, which ultimately culminate in reduced brain size. Notably, our findings align with emerging evidence highlighting the critical role of mitotic regulation in neurodevelopment, yet they also expand the known genetic landscape of microcephaly. Compared with previous studies focusing on single-gene-protein interactions, our identification of the AURKA-NUSAP1-KIF2C pathway provides a more integrated view of the molecular mechanisms underlying this disorder.

## Supporting information

Supplemental Figure 1, Figure2 and Supplemental Table1, and will be used for the linkl to the file on the preprint site.

## Acknowledgement

This project is supported by the National Key Research and Development Program (2024YFC2707002, 2023YFC3403000), Innovation Program of Shanghai Municipal Education Commission (2023ZKZD16), National Natural Science Foundation of China (82071262, 32300464), Natural Science Foundation of Hunan Province (2023JJ30332), Shanghai Municipal Science and Technology Major Project (20JC1418600), China Postdoctoral Science Foundation (2023M732266), Key Technology Breakthrough Program of Ningbo Sci-Tech Innovation YONGJIANG 2035 (2024Z221), Municipal Public Welfare Project (2022S035), Ruixin Project of Hunan Provincial Maternal and Child Health Care Hospital (2023RX01), Major Scientific and Technological Projects for Collaborative Prevention and Control of Birth Defects in Hunan Province and Shanghai Jiao Tong University STAR Grant (YG2023ZD26, YG2023LC14).

The last author would like to thank Dr. Tan Guanhao and Dr. Li Zongju for their unwavering support and unconditional love throughout the research journey.

