## Supplemental Figure 1, Figure2 and Supplemental Table1, and will be used for the linkl to the file on the preprint site. for "NUSAP1 regulates mitotic processes via KIF2C interaction and AURKA phosphorylation in primary microcephaly"

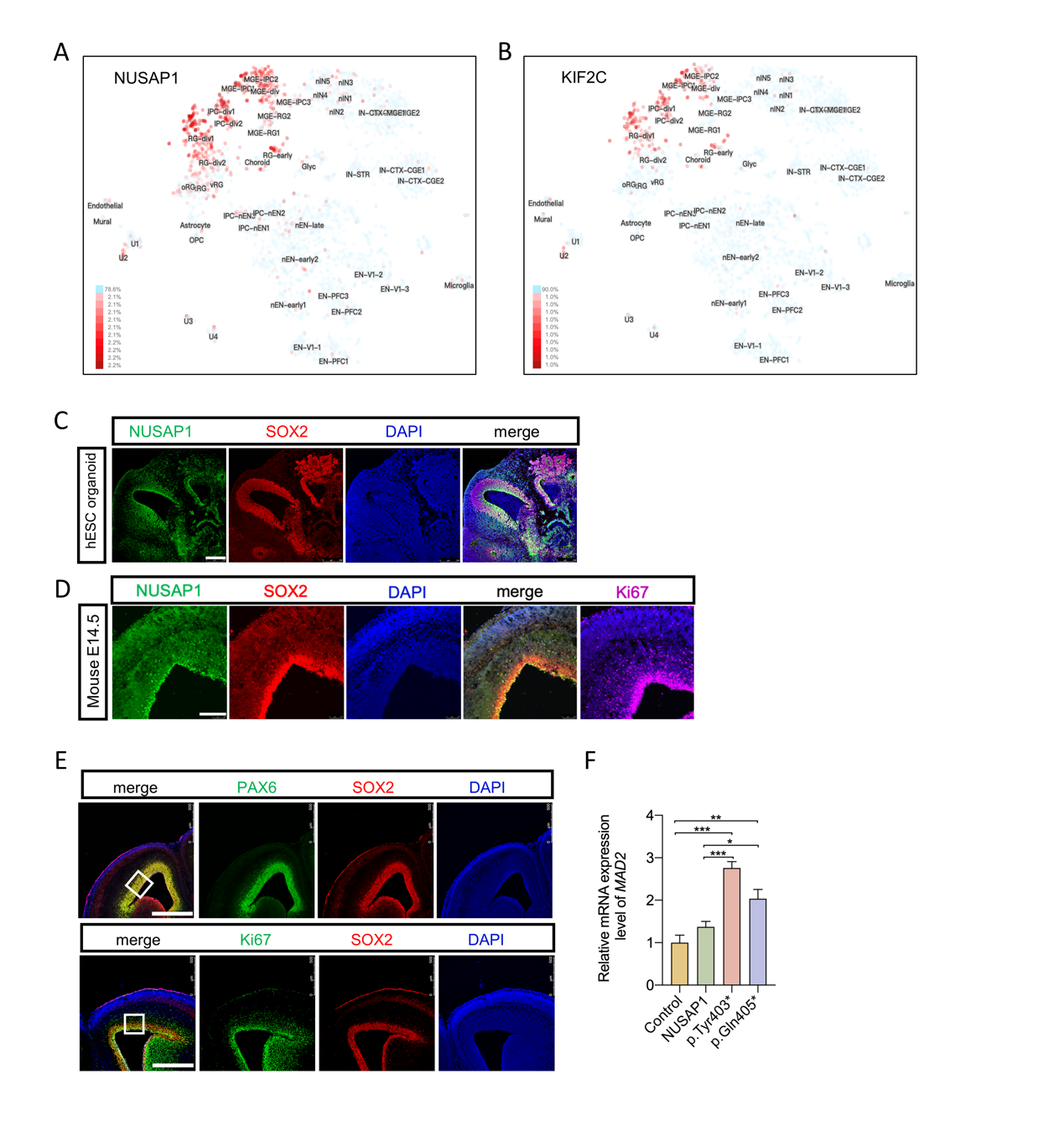

**Figure S1. NUSAP1 and KIF2C are specifically expressed in NSPCs. (A)** NUSAP1 expression and distribution in developing human fetal neocortex (https://cells.ucsc.edu). (**B)** KIF2C expression and distribution in developing human fetal neocortex (https://cells.ucsc.edu). (**C)** Immunofluorescent images of the hESC organoid stained for NUSAP1 (green) and SOX2 (red) were used to detect the co-expression and co-localization of these two proteins. DAPI was used to stain the nuclei (blue). Merged images of the NUSAP1, KIF2C, and DAPI channels are shown on the right. Scale bar = 250 μm. **(D)** Immunofluorescent images of the E14.5 mouse cerebral cortex stained for NUSAP1 (green), SOX2 (red) were used to detect the co-expression and co-localization of these two proteins. Ki67 was used to mark the proliferating cells. DAPI was used to stain the nuclei (blue). Scale bar = 200 μm. **(E)** Immunofluorescent images of the cerebral cortex stained harvested at E14.5 for nuclei (DAPI - blue), SOX2 (red), PAX6 (upper panel, green), and Ki67 (lower panel, green). The merged image was shown on the left. Scale bar = 500 μm. The white squares represented the location of the cerebral cortex that were selected for the Figure 2 images. (**F)** The relative mRNA expressions of *BUBR1* was calculated by comparing their expression values with those in control cells, which were normalized to 1. Unpaired two-tailed t-test was used to compare different experimental groups for significant differences. Data are shown as mean±SEM. **P* < 0.05, ***P* < 0.01, ****P* < 0.001.

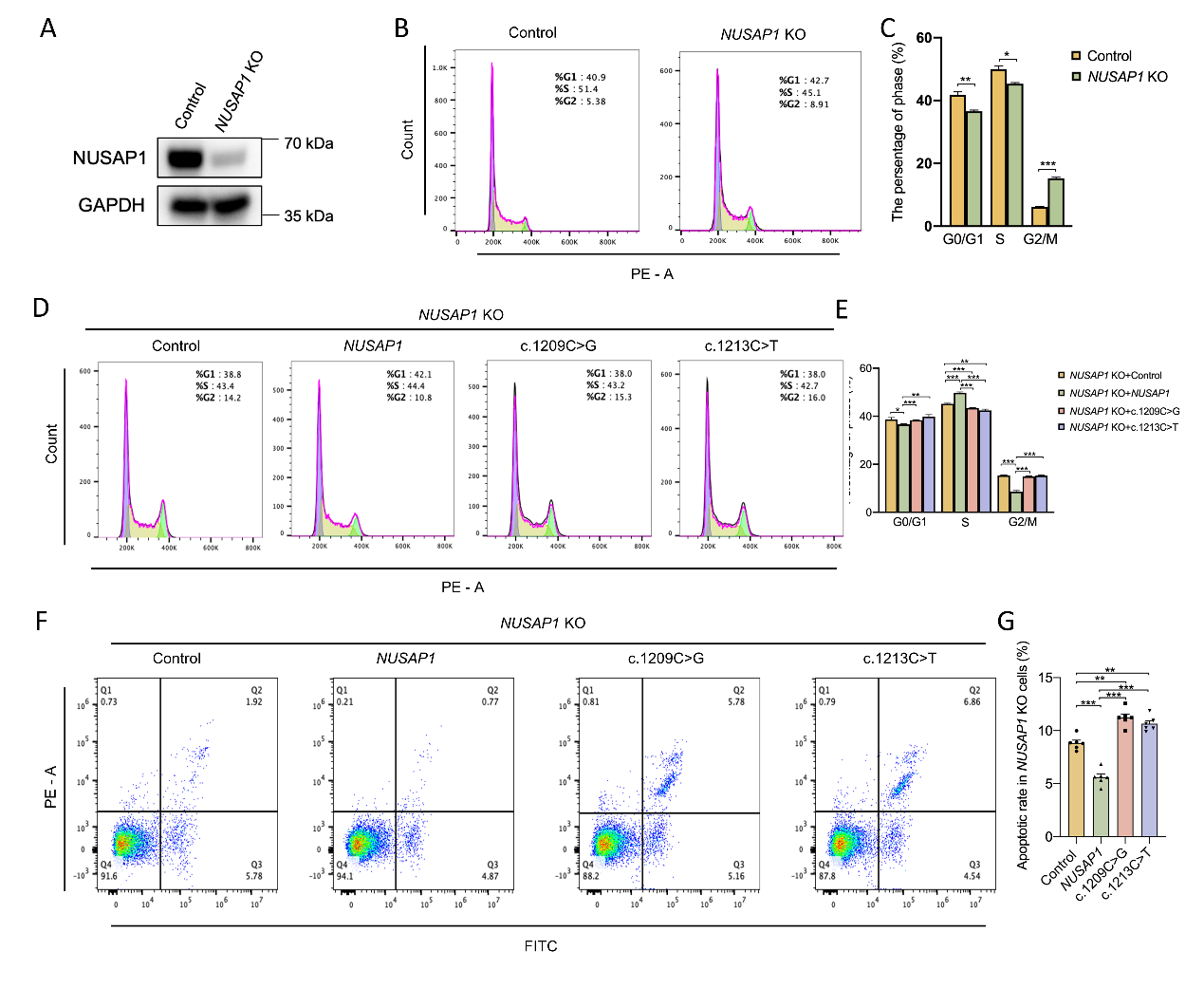

**Figure S2. Failure of NUSAP1 mutant forms to rescue knockout-induced phenotypes in HEK293T cells. (A)** *NUSAP1* knockout efficiency of HEK293T cells was analyzed by Western blotting. **(B, C)** Cell cycle was analyzed in *NUSAP1* KO cells by flow cytometry (G0/G1, S and G2/M phases). **(D, E)** Cell cycle was analyzed by flow cytometry in *NUSAP1* KO HEK293T cells transfected with control, *NUSAP1*, c.1209C>G or c.1213C>T plasmids, showing the distribution of cells in G0/G1, S and G2/M phases. **(F, G).** Apoptosis was analyzed by flow cytometry in *NUSAP1* KO HEK293T cells transfected with control, *NUSAP1*, c.1209C>G or c.1213C>T plasmid. Unpaired two-tailed t-test was used to compare different experimental groups for significant differences. Data are shown as mean±SEM. **P* < 0.05, ***P* < 0.01, ****P* < 0.001.

Table S1. Clinical features.

|  |  | **Patient 1 (Family_1)** |  | **Patient 2 (Family_2)** |
| --- | --- | --- | --- | --- |
| Age at last evaluation |  | 7 years and 5 months |  | 5 years and 6 months |
| Gender |  | Male |  | Male |
| Prenatal ultrasound |  | Microcephaly  (21.2 cm -3SD at 26 weeks; 24.7 cm -3.3SD at 30 weeks; 28.6 cm -3.3SD at 36 weeks) |  | Microcephaly |
| Family medical history |  | No |  | No |
| Gestational age |  | Full-term |  | - |
| Height and weight |  | normal |  | normal |
| Head circumference |  | 30 cm (-3.5 SD) at birth;  44 cm (-6 SD) at 7 years and 5 months |  | 31.5 cm (-3.4SD) at 29 days old;  45.7 cm (-4.1SD) at 5 years and 6 months |
| Developmental delay |  | Severe |  | Severe |
| Epilepsy |  | Began on 17 days old  (eyelid fluttering, ocular deviation, head turning, upward gaze, absence seizures, and generalized tonic seizures) |  | Began on 29 days old  (focal seizures that progressed to secondary generalization) |
| Electroencephalogram (EEG) |  | Interictal right-sided sharp waves and sharp-slow wave complexes. Ictal recordings revealed rhythmic right-sided spike-slow wave discharges |  | Multifocal epileptiform activity, including spikes and sharp-slow waves |
| Brain MRI |  | Partial agenesis of the corpus callosum |  | Cerebral dysgenesis, pachygyria, thinning of the corpus callosum, and ventriculomegaly |
| Vocal disturbances |  | Severe (lacked even basic speech) |  | Severe (lacking any meaningful speech) |
| Feeding |  | Maintained on an exclusive liquid diet. |  | - |
| Motor functions |  | Attaining only basic motor functions |  | Unable to sit, crawl, or roll |
| Others |  | Generalized hypertonia and emerging signs of precocious puberty |  | Left side cryptorchidism |
| Exome sequencing |  | *NUSAP1* (NM_016359.4): c.1213C>T; p.Gln405* |  | *NUSAP1* (NM_016359.4): c.1209C>G; p.Tyr403* |

The symbol “-” indicates that the relevant records were unavailable.
